# Coordination between metabolic transitions and gene expression by NAD^+^ availability during adipogenic differentiation in human cells

**DOI:** 10.1101/2021.10.04.462470

**Authors:** Edgar Sanchez-Ramírez, Thi Phuong Lien Ung, Ximena del Toro-Rios, Guadalupe R. Fajardo-Orduña, Lilia G. Noriega, Armando R. Tovar, Juan José Montesinos, Ricardo Orozco-Solís, Chiara Stringari, Lorena Aguilar-Arnal

## Abstract

Adipocytes are the main cell type in adipose tissue, a critical regulator of metabolism, highly specialized in storing energy as fat. Adipocytes differentiate from multipotent mesenchymal stromal cells through adipogenesis, a tightly controlled differentiation process involving closely interplay between metabolic transitions and sequential programs of gene expression. However, the specific gears driving this interplay remain largely obscure. Additionally, the metabolite nicotinamide adenine dinucleotide (NAD^+^) is becoming increasingly recognized as a regulator of lipid metabolism, being postulated as promising therapeutic target for dyslipidemia and obesity. Here, we explored the effect of manipulating NAD^+^ bioavailability during adipogenic differentiation from human mesenchymal stem cells. We found a previously unappreciated strong repressive role for NAD^+^ on adipocyte commitment, while a functional NAD^+^-dependent deacetylase SIRT1 appeared crucial for terminal differentiation of pre-adipocytes. Remarkably, repressing the NAD^+^ biosynthetic salvage pathway during adipogenesis promoted the adipogenic transcriptional program, suggesting that SIRT1 activity during adipogenesis is independent from the NAD^+^ salvage pathway, while two photon microscopy and extracellular flux analyses suggest that its activation relies on the metabolic switch. Interestingly, SIRT1-directed control of subcellular compartmentalization of redox metabolism during adipogenesis was evidenced by two-photon fluorescence lifetime microscopy.

**Significance Statement:** Adipocyte differentiation occurs from mesenchymal stem cells through the adipogenic process, involving sequential activation of both transcriptional and metabolic programs in a tightly coordinated manner. However, how transcriptional and metabolic transitions reciprocally interact during adipogenic differentiation remains largely obscure. Here we describe that the metabolite NAD^+^ is suppresses adipogenesis trough rewiring transcription, while a functional NAD^+^-dependent deacetylase SIRT1 is essential for terminal differentiation of pre-adipocytes. Using two-photon fluorescence lifetime microscopy, we created a metabolic map of NADH and lipid content simultaneously in live cells and described a new role for SIRT1 in the control of compartmentalization of redox metabolism during adipogenesis. These findings advance our understanding to improve therapeutical approaches targeting the NAD^+^-SIRT1 axis as treatment for obesity and dyslipemia.

## INTRODUCTION

Adipose tissue is a crucial regulator of body metabolism through storing calories as lipids in response to excessive nutritional intake and serving as a source of energy by mobilizing these lipids during starvation, amongst others. Notably, the adipose tissue is a relevant endocrine organ, producing several adipokines such as leptin or adiponectin(1). It is primarily composed of adipocytes, and a fraction of a heterogeneous collection of cell types which include mesenchymal stem cells (MSC), endothelial precursors, immune cells, smooth muscle cells, pericytes and preadipocytes(2). Disfunction of the adipose compartment is common to metabolic diseases including obesity or type 2 diabetes. Indeed, increased white adipose tissue (WAT) mass observed in obesity is due to both adipocyte hypertrophy and increased proliferation and differentiation of adipocyte progenitors(3, 4), which originate from MSCs though the adipogenic process (5, 6). Hence, understanding the mechanisms underlaying adipogenesis in humans is crucial to design therapeutic strategies for prevalent metabolic dysfunctions.

MSCs are multipotent progenitor cells able to differentiate to osteoblasts, myocytes, chondrocytes, and adipocytes. Fate decision is determined by specific signaling pathways such as transforming growth factor-beta (TGFβ)/bone morphogenic protein (BMP) signaling, wingless-type MMTV integration site (Wnt) signaling or fibroblast growth factors (FGFs)(7, 8). In particular, the adipogenic process occurs in two major phases: commitment to progenitors and terminal differentiation; both of which are tightly regulated by intertwined transcriptional, epigenomic and metabolic transitions(8). At the transcriptional level, the master regulators of adipogenesis are the key transcription factors peroxisome proliferator-activated receptor γ (PPARγ) and CCAAT/enhancer binding protein α (C/EBPα)(9, 10), which promote growth arrest and the progressive expression of a lipogenic transcriptional program including the hormones adiponectin and leptin, and the lipases adipose triglyceride lipase (ATGL) and lipoprotein lipase (LPL)(8). Concomitantly, a switch from highly glycolytic to oxidative metabolism with increased mitochondrial reactive oxygen species (ROS) is essential for adipocyte differentiation(11, 12). However, how transcriptional and metabolic transitions reciprocally interact during adipogenic differentiation remains an open question.

During the past few years, energy metabolism is becoming increasingly recognized as an effective therapeutic target for obesity. Specifically, therapies aiming to increase endogenous nicotinamide adenine dinucleotide (NAD^+^) levels have been proven effective to reduce adiposity in both mouse and human(13–16). Indeed, NAD^+^ levels decline in metabolic tissues of obese mice and humans(14, 15, 17–20), which may contribute to metabolic disfunction by, for example, reducing the activity of SIRT1, a deacetylase using NAD^+^ as cofactor and known to regulate mitochondrial function and metabolism(21, 22). These evidences suggest that NAD^+^ metabolism might be a central player on adipose tissue homeostasis probably by regulating mitochondrial function and consequently, adipocyte differentiation. Along these lines, in mouse preadipocytes, NAD^+^ synthesis through the salvage pathway and SIRT1 activity appear essential for adipogenesis(23); however, the interplay between NAD^+^ bioavailability and SIRT1 function during adipogenesis in humans remains poorly understood.

In this study, we explored the coordinated dynamics of the transcriptional and energy metabolism reprogramming during adipogenic differentiation of human MSC (hMSC). Using two-photon fluorescence lifetime microscopy (2P-FLIM) on live hMSC, we created a non-invasive metabolic map of NADH compartmentalization at a submicron resolution to define the dynamics of redox metabolism during adipogenesis, which appeared tightly synchronized with mitochondrial function and transcriptional reprogramming. Moreover, we described a previously unappreciated robust inhibitory role for NAD^+^ on adipocyte commitment, while SIRT1 activity appeared essential for terminal differentiation. Surprisingly, suppressing the NAD^+^ salvage pathway during adipogenesis led to increased expression of adipocyte markers, indicating that SIRT1 activation during adipogenesis doesńt depend on NAD^+^ biosynthesis through the salvage pathway. Remarkably, SIRT1-directed control of compartmentalization of redox metabolism during adipogenesis was evidenced by 2P-FLIM.

## RESULTS

### NAD^+^ obstructs adipogenic differentiation and lipid accumulation in hMSC

To approach the question whether variations in NAD^+^ bioavailability impact adipogenesis, we induced adipogenic differentiation in hMSCs in the presence or absence of NAD^+^ (Figure 1A). Interestingly, NAD^+^ treatment obstructed neutral lipid accumulation, as visualized by significantly less accumulation of Oil-red-O (ORO) stain at terminal differentiation than in non-treated (NT) cells (Figure 1A, day 16, *P*<0,01, Two-way ANOVA with Tukey’s post-test). Concomitantly, a treatment with FK866, a potent and highly selective inhibitor of NAMPT(24, 25), the rate limiting enzyme in the NAD^+^ salvage biosynthetic pathway, led to significantly more neutral lipids at day 16 (Figure 1A, day 16, *P*<0,01, Two-way ANOVA with Tukey’s post-test), further reinforcing the idea that NAD^+^ bioavailability might oppose lipogenesis. As it has been largely shown that SIRT1 is a mayor effector of NAD^+^ signaling, we reasoned that it might be dispensable for adipogenesis. Surprisingly, selective inhibition of SIRT1 during adipogenic differentiation strongly hindered neutral lipid accumulation (Figure 1A, day 16, P<0,01, Two-way ANOVA with Tukey’s post-test). Interestingly, distinct dynamic changes in lipid accumulation were observed at days 4, 8 and 12 of adipogenic inductions with the different treatments (Figure 1A).

**Figure 1:**
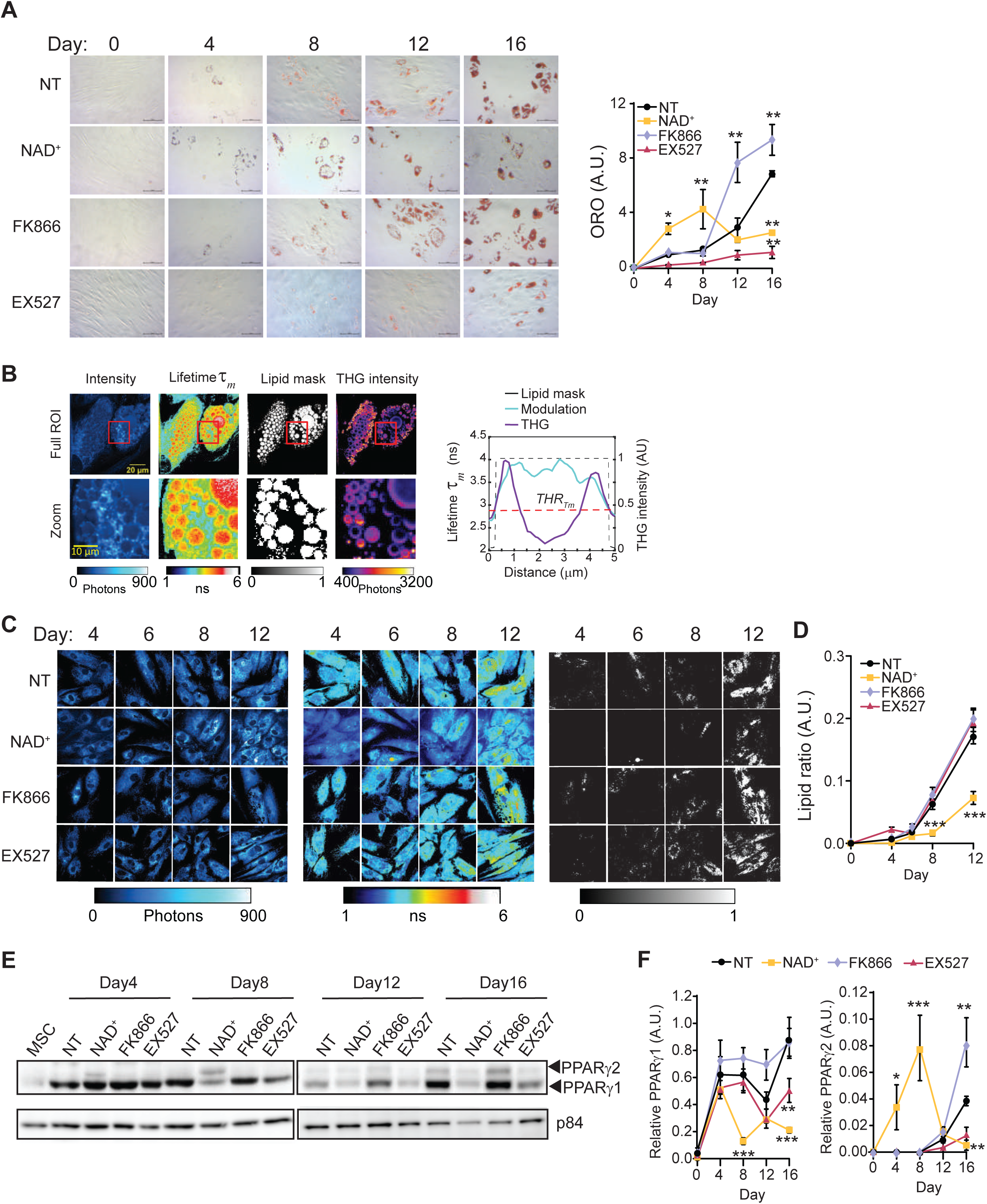
NAD^+^ hinders adipogenic differentiation and lipid accumulation in hMSC. Adipogenic differentiation was induced in hMSC in the absence or presence of the indicated drugs: NAD^+^ (5 mM), FK866 (1 nM) or the SIRT1-specific inhibitor EX527 (50 μM). **A)** Neutral lipids were stained with Oil-Red-O (ORO) at the indicated days after adipogenic induction, and representative images from are shown. Quantification was performed by densitometry from n=4 technical and 3 biological replicates. **B)** Representative images from Label free quantification of lipid droplets by Fluorescence Lifetime Microscopy of an adipocyte at terminal differentiation. Intensity , modulation lifetime tauM (τ_m_) lipid mask determined by a τ_m_ threshold, and Third harmonic generation (THG) intensity are shown as indicated. The graph on the right shows a cross sectional profile of lipid mask created by a τ_m_ threshold and THG signal of one lipid droplet of 5 µm diameter. **C)** Representative images of intensity (left), modulation lifetime τ_m_ (middle) and lipid mask (right) of hMSCs during adipogenic differentiation at the indicated days of culture under selected treatments **D)** Quantification from lipid ratio at indicated days and treatments during the differentiation process of hMSC. n=14 single cells; experiments were conducted in triplicate. **E)** PPARγ1 and PPARγ2 protein expression levels were measured by western blot in whole cell extracts at the indicated days after adipogenic induction. p84 was used as loading control. **F)** quantification of PPARγ1 and PPARγ2 protein expression, normalized to p84 loading control. n=3 biological replicates. For all graphs, data is presented by mean ± SEM (*p< 0,05, **p< 0,01, ***p< 0,001, Two-way ANOVA with Tukey’s post-test)

To further confirm that lipid accumulation is changing with the treatments in living cells while avoiding potential artifacts from fixation or ORO stain, we performed label-free Fluorescence lifetime microscopy of intrinsic lipid-associated fluorophores in living cells (see Methods section). Third harmonic generation (THG) images were used as reference to identify lipid droplets (26, 27) (Figure 1B, S1A, S1B), and they showed bright round dots for lipid droplets smaller than 1µm and hollow round structures at the interface between cytoplasm and lipid droplets larger than 1µm. FLIM images (Figure 1B) indicated that lipid droplets associated fluorophores have a longer lifetime (τ_m_) with respect to the rest of the cell cytoplasm, hence we determined a threshold to automatically and systematically identify and segment lipid droplets (Figure 1B, S1B, Methods section). This method allowed us to quantitively assess lipid accumulation in live hMSCs during adipogenic induction (Figure 1C). We imagined cells at days 4, 6, 8 and 12, corresponding to those when more dynamic changes were previously observed, and found that NAD^+^ treatment significatively impaired lipid accumulation after day 8 (Figure 1C, D; day 8 and 12, *P*<0.001, Two way ANOVA with Tukey’s post-test). With this approach, we didńt find significant differences in lipid accumulation in live cells when comparing FK866 and EX527 treatments to non-treated cells.

PPARγ is considered a master transcription factor for adipogenesis(28, 29). It′s expression is induced in early stages(30), and sustains the specific transcriptional program for adipocyte differentiation(31). For these reasons, we explored how NAD^+^ levels impact PPARγ protein expression during adipogenic induction of hMSCs (Figure 1E, 1F). We found that PPARγ1 isoform is strongly expressed since day 4 of differentiation for all tested conditions (Figure 1E, F). Yet, while its expression was overall sustained along the adipogenic process in control cells, NAD^+^ triggers a very significant reduction in PPARγ1 protein expression while favoring expression of the PPARγ2 isoform at differentiation day 8, which is lost at later stages (Figure 1E, F; *P*<0.01, Two-way ANOVA with Tukey’s post-test). Interestingly, EX527 treatment led to significantly decreased expression of PPARγ1 at the end of differentiation. Conversely, FK866 significantly increased PPARγ2 isoform with respect to the control without affecting PPARγ1 protein expression. As expected, a 16-days treatment of hMSC with any of the compounds led to detectable changes in PPARγ expression (Figure S1C). Together, these differential dynamics on PPARγ protein expression point to a mayor role for transcriptional control in the adipogenic potential shown after each treatment, which is also in line with the distinct outcomes in lipid accumulation.

### Extensive and specific reprogramming of the transcriptome during adipogenic differentiation by NAD^+^

To decipher the dynamic changes occurring in the transcriptome of hMSC during differentiation, we performed RNA-seq analyses from undifferentiated hMSC, and differentiated cells at two different time points: at 8 days of adipogenic differentiation (middle of differentiation protocol) and at 16 days (end of differentiation), in untreated cells and in cells treated with NAD^+^, FK866 or EX527. We first performed an unbiased principal component analysis (PCA) revealing that, as expected, the largest variation was due to the differentiation process (PC1, 24,92%; Figure 2A and S2A). Interestingly, the second component retaining 21.03% of the original variance between samples was mostly related to NAD^+^ treatment (Figure 2A, S2A). Finally, a third component could be due to the progress of the differentiation process itself (PC3, 8.98%; Figure 2A, S2A). We then performed differential gene expression analyses to compare between samples using DESeq2 (32), (see methods for details and Table S1 for normalized counts). These analyses further reinforced that NAD^+^ treatment had a large impact in the transcriptome, comparable to the differentiation process itself, as more than 3000 genes were differentially expressed (DE) in differentiating cells treated with NAD^+^ when compared with their untreated controls (adjusted *P* <0.05) (Figure S2B). Concomitantly, inhibition of NAD^+^ biosynthesis by FK866 lead to just 171 differentially expressed genes when compared with the untreated controls at the end of the adipogenic process (Figure S2B, AD16_FKvsAD16). Hence, we sought to determine the molecular signatures of NAD^+^ treatment during adipogenic differentiation of hMSC. To do so, we first selected the transcripts specifically dysregulated by NAD^+^ treatment and found 660 common DE genes in NAD^+^ treated cells when compared with any other treatment or hMSC, which conform a unique NAD^+^ molecular signature (Figure 2B). Out of these, 157 genes were consistently upregulated, while 407 were always silenced by the treatment (Figure 2B, Table S2). Next, we sought to explore the molecular routes responsible for the impaired adipogenic capacity of NAD^+^ treated cells. Hence, we selected the genes which appeared consistently dysregulated in NAD^+^ treated cells whAmong these, en compared with the rest of differentiating cells (AD, AD_EX, AD_FK). We identified 2,057 of these genes at day 8 and 1,969 at day 16 after adipogenic induction. 993 genes were shared, out of which 70% (703) were consistently downregulated, while 28% (279) were always overexpressed (Figure 2C, 2D, Table S2). Gene ontology (GO) analyses revealed that most of the upregulated genes are implicated in apoptotic and response to stress processes, while downregulated genes relate to mRNA metabolism, cellular motility, and differentiation processes (Figure 2E, 2F, Table S2). A subsequent pathway mapping in KEGG of the same transcripts revealed that many genes involved in steroid biosynthesis (*SQLE, DHCR7, FDFT1, SC5D, MSMO1*) were upregulated by NAD^+^ treatment during adipogenic differentiation of hMSCs. Also, apoptotic signaling pathways appeared active, as suggested by the overexpression of several death receptors (*TNFRSF -10A*, -*10B*, -*10D*) , probably triggered by activated ER stress and unfolded protein response, evidenced by increased transcription of *IRE1α* (*ERN1*), BiP (*HSPA5*) and *XBP1* (Fig. 1G, Table S2). Interestingly, increased cholesterol biosynthesis induces ER stress in macrophages(33). In contrast, a unique set of genes downregulated by an excess of NAD^+^ during adipogenic differentiation pertained to the ribosome pathway, involving many transcripts for ribosomal proteins (Figure 2H, S3A), indicating that their expression is strongly suppressed by NAD^+^ in an adipogenic context. Additionally, many transcripts involved in cell adhesion and motility, which become expressed during differentiation, were downregulated in NAD^+^ treated cells, indicating that the adipogenesis process is arrested. Furthermore, we found that the JAK-STAT pathway was impaired in NAD^+^ treated cells, with downregulated transcripts including *STAT5A and STAT5B,* known positive regulators of the master adipogenic TF PPARγ (34), or the leptin encoding gene *LEP* (Figure 2H, Table S2). Taken together, these data points towards NAD^+^ promoting a proapoptotic and anti-adipogenic environment, therefore inhibiting differentiation and maturation of adipocytes. Accordingly, a promoter screening for transcription factor binding motifs revealed significant enrichment for SMAD motifs within promoters of upregulated genes (Figure 2I, *P* = 10^-7^), and for CEBP:AP1 motifs amongst downregulated geneś promoters (Figure 2J, *P* = 10^-8^). Indeed, SMADS are well known apoptotic regulators, while CEBP is a master adipogenic TF(29, 35).

**Figure 2:**
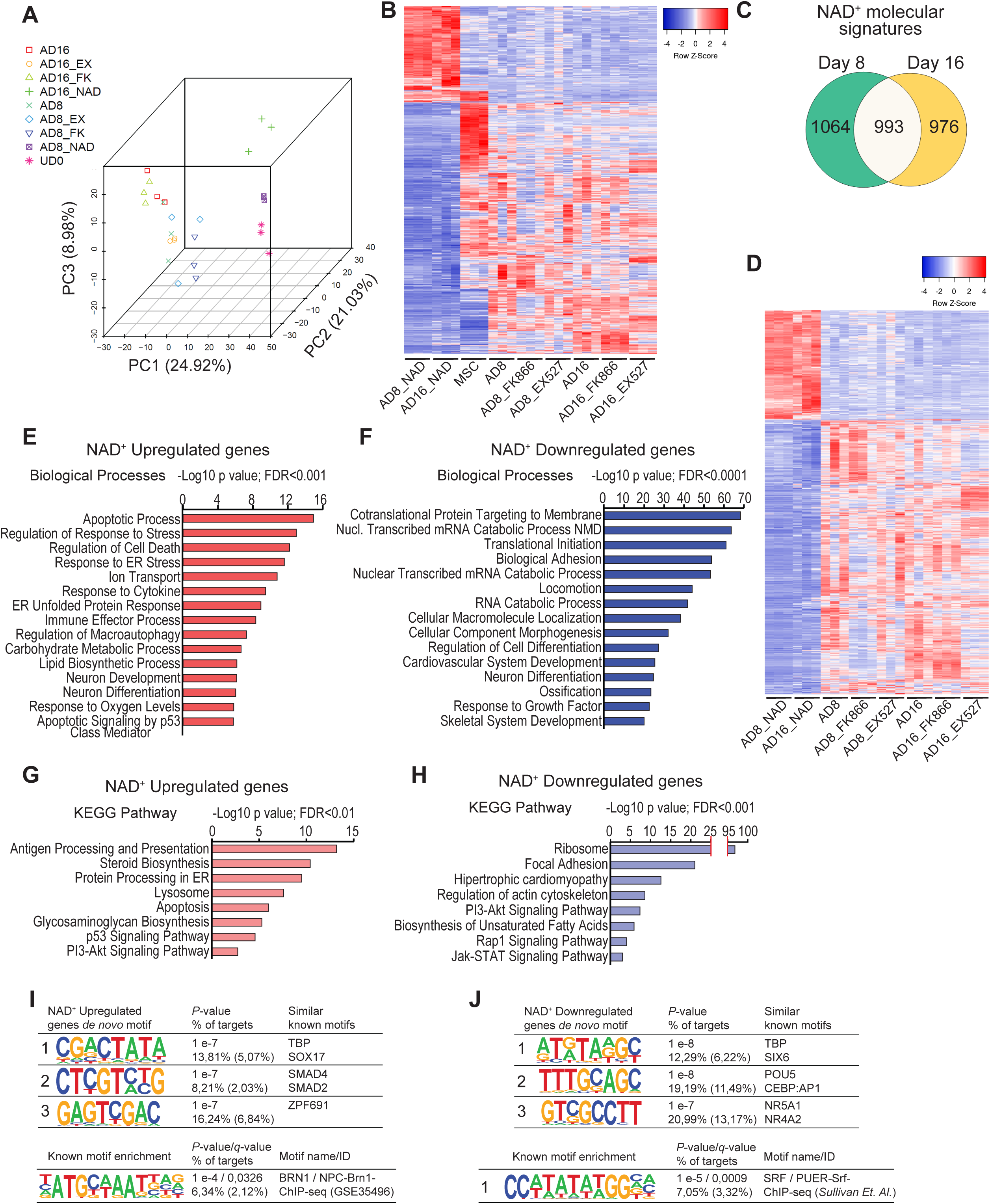
NAD^+^ treatment during adipogenesis from hMSC induces profound and specific changes in the transcriptome. RNA-seq was performed per triplicate from multipotent hMSC (MSC, UD0), or at days 8 (AD8) and 16 (AD16) after adipogenic induction in the absence (AD8, AD16) or presence of 5 mM NAD+ (AD8_NAD, AD16_NAD), 1 nM FK866 (AD8_FK866, AD16_FK866) or 50 μM EX527 (AD8_EX527, AD16_EX527) **A)** Principal component analysis (PCA) was computed for the whole data. **B)** Heatmap comparing expression from 660 genes DE exclusively in NAD^+^ treated cells (FDR-adjusted *P*-value <0.05) **C)** Overlap of DE transcripts between NAD+ treated cells and the rest of the tested conditions at day 8 and day 16 after adipogenic induction **D)** Heatmap comparing expression from 994 genes DE exclusively in NAD^+^ treated cells at both days 8 and 16 after adipogenic induction when compared with the rest of the samples. **E-H)** Functional annotation of the 994 DE genes constituting the NAD^+^ transcriptional signature: biological processes (E, F) or KEGG pthaways (G-H) for consistently upregulated (E, G) or downregulated (F, H) transcripts. **I, J)** Homer *de novo* motif discovery analyses from promoters of genes specifically upregulated (I) or downregulated (J) after NAD^+^ treatment during adipogenic induction.

### SIRT1 regulates terminal differentiation of pre-adipocytes but is dispensable for adipogenic commitment

If an excess of NAD^+^ hinders the adipogenic transcriptional reprogramming, while inhibiting NAD^+^ biosynthesis has only a mild transcriptional effect at the end of the adipogenic process, it is conceptually apparent that critical NAD^+^ consuming enzymes such as SIRT1 would not be essential for adipocyte differentiation. Accordingly, SIRT1 inhibition by EX527 had only a mild effect on the transcriptome at day 8, with just 57 DE genes compared with the untreated cells (Figure S2B, AD8_EX vs AD8, Table S3). Strikingly, in terminally differentiated adipocytes we found 2040 differentially expressed genes, with 1095 upregulated and 945 downregulated transcripts (Figure S2B, AD16_EX vs AD16, Figure 3A, Table S3). This is in line with the absence of lipid accumulation in these cells, and indicates that probably, SIRT1 is dispensable for lineage commitment, but essential for terminal differentiation of adipocytes. Interestingly, at day 8 of differentiation, the anti-adipogenic genes *EGR1* and *NR4A1* (36, 37) were overexpressed in EX527 treated cells (Table S3, *P*<0,0001, Fold change > 2), and these have previously been shown to be upregulated in the early mitotic clonal expansion phase during preadipocyte differentiation(37), further reinforcing that SIRT1 inhibition compromises maturation, but not commitment of preadipocytes. Accordingly, at day 16 many genes pertaining to pathways such as fatty acid metabolism, degradation or lipolysis were downregulated in cells treated with EX527, most of them implicated in PPAR signaling (Figure 3B, S4A, Table S3). Amongst these, the adipokine genes leptin (*LEP*), leptin receptor (*LEPR*), and adiponectin (*ADIPOQ*) showed significatively lower levels in EX527 treated cells (Table S3). Interestingly, SIRT1 inhibition upregulated transcripts enriched for adhesion and locomotion, critical processes during the onset of the adipogenesis, including the fibronectin gene *FN1*, which inhibits adipocyte maturation(38) (Figure 3C, S4B. Table S3). Accordingly, gene set enrichment analysis (GSEA) identified adipogenesis and apical junction as the top-ranked hallmarks enriched in untreated and EX527 treated cells respectively (Figure 3D, E). Together, these data indicate that SIRT1 activity is essential for adipogenic differentiation, specifically by tightly controlling the transition between adipocyte commitment and maturation, and the maturation process itself. Remarkably, even though SIRT1 mRNA expression increased at the beginning of adipogenic induction (Figure 3F), the protein significantly increased after 8 days of adipogenic induction (Figure 3G, *P*<0,0001, Kluskal-Wallis with Dunńs post-test), probably due to posttranscriptional regulation, and further reinforcing the notion that SIRT1 is essential for adipocyte maturation. Strikingly, a motif analysis revealed that pharmacological inhibition of SIRT1 downregulated preferred targets genes for distinct members of the FOX family of transcription factors (Figure 3H, *P*=1E-11), while overexpressed genes were enriched for motifs binding homeobox (NKX3-1,-2) or b-Helix-loop-helix (b-HLH) transcription factors such as CLOCK:BMAL (Figure 3I, *P*=1E-8). Indeed, most of these are well-known targets for SIRT1 deacetylation.

**Figure 3:**
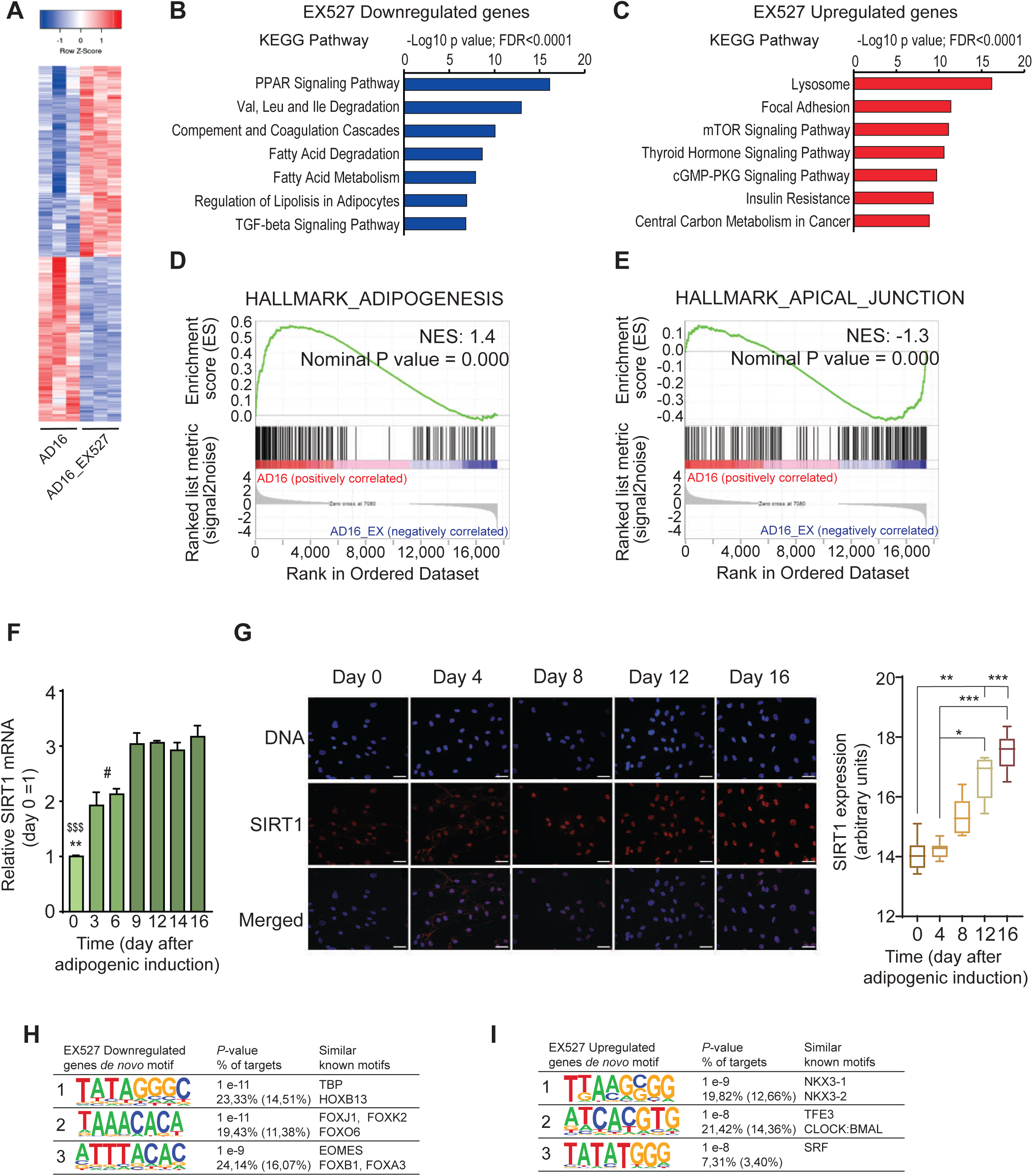
SIRT1 activity is essential for terminal differentiation of pre-adipocytes. **A)** Heatmap comparing expression from 2040 genes DE between EX527-treated cells (50 μM) and untreated cell at day 16 after adipogenic induction on hMSC. (FDR- adjusted *P*-value <0.05) **B, C)** KEGG pathway enrichment analyses from genes downregulated (B) or upregulated (C) by EX527 treatment during adipogenic differentiation, at day 16 after induction, compared with untreated, terminally differentiated adipocytes. **D, E**) Gene set enrichment analysis (GSEA) investigated within the molecular signature database (MSigDB) “Hallmark” gene set collection. Genes were rank-ordered by differential expression between terminally differentiated adipocytes untreated (AD16) or treated with EX527 (AD16_EX). **F)** *SIRT1* gene expression levels were assessed by RT-qPCR at the indicated days after adipogenic induction on hMSC. n= 3 biological and 2 technical replicates. One-way ANOVA followed by Tukeýs post test. * p <0.05, ** p <0.01, *** p <0.001. Symbol key for multiple comparisons: *: day 0 vs days 3, 6; $: day 0 vs days 9-16; #: days 3, 6 vs days 9-16. **G)** SIRT1 protein expression and subcellular location was explored by immunofluorescence at the indicated days after adipogenic induction on hMSC. Boxplot shows densitometric analyses from n= 2 biological and 7 technical replicates. Kluskal-Wallis test followed by Dunn’s multiple comparisons test was applied. * p <0.05, ** p <0.01, *** p <0.001. **H, I)** Homer *de novo* motif discovery analyses from promoters of genes downregulated (H) or upregulated (I) by EX527 treatment during adipogenic induction, at terminal differentiation (day 16).

SIRT1 activity heavily relies on NAD^+^ availability, hence the treatment with the NAMPT inhibitor FK866, known to dampen intracellular NAD^+^ levels, would prevent SIRT1 activation and the subsequent adipocytic gene expression program. Surprisingly, FK866 treatment during adipogenesis led to enhanced adipocytic differentiation, through overexpression of adipogenic hallmarks and increased adipocytokine and PPAR signaling over the control, non-treated adipocytes, at terminal differentiation (Figure 4A,B, Table S4). Indeed, 39 genes involved in lipid metabolism, including *CD36*, *CEBPA*, *ADIPOQ*, *LPL*, *ACLY*, *FASN*, *ACACA*, *ACACB*, *PLIN1* and *RETSAT*, were highly expressed in terminal adipocytes treated with FK866 (Figure 4C). Moreover, genes related to ossification or cartilage development were downregulated in these cells (Figure 4D), indicating that low NAD^+^ levels induced by FK866 treatment potentiate the adipogenic over the osteogenic lineage in hMSCs(7). This is in line with our previous observation that NAD^+^ treatment hinders differentiation of hMSC to adipocytes; yet, it is in contrast with the need of SIRT1 activity, which relays on its cofactor NAD^+^, for adipocyte maturation. Furthermore, out of 57 differentially expressed genes between the control and EX527 treated cells at day 8 of differentiation, 33 (58%) were also dysregulated by FK866 treatment, suggesting that SIRT1 is not active in FK866 treated cells at day 8 (Figure 4E). Accordingly, SIRT1 expression both at the mRNA and protein levels was dampened by EX527 and FK866 to similar levels during adipogenic induction (Figure 4F, Two-way ANOVA with Turkeýs post-test; Figure 4G, Kluskal-Wallis with Dunńs post-test; Figure S5A). These data suggest that SIRT1 function might be dispensable for adipocyte commitment, but necessary for differentiation, and the source of NAD^+^ essential for SIRT1 activity does not require the salvage pathway.

**Figure 4:**
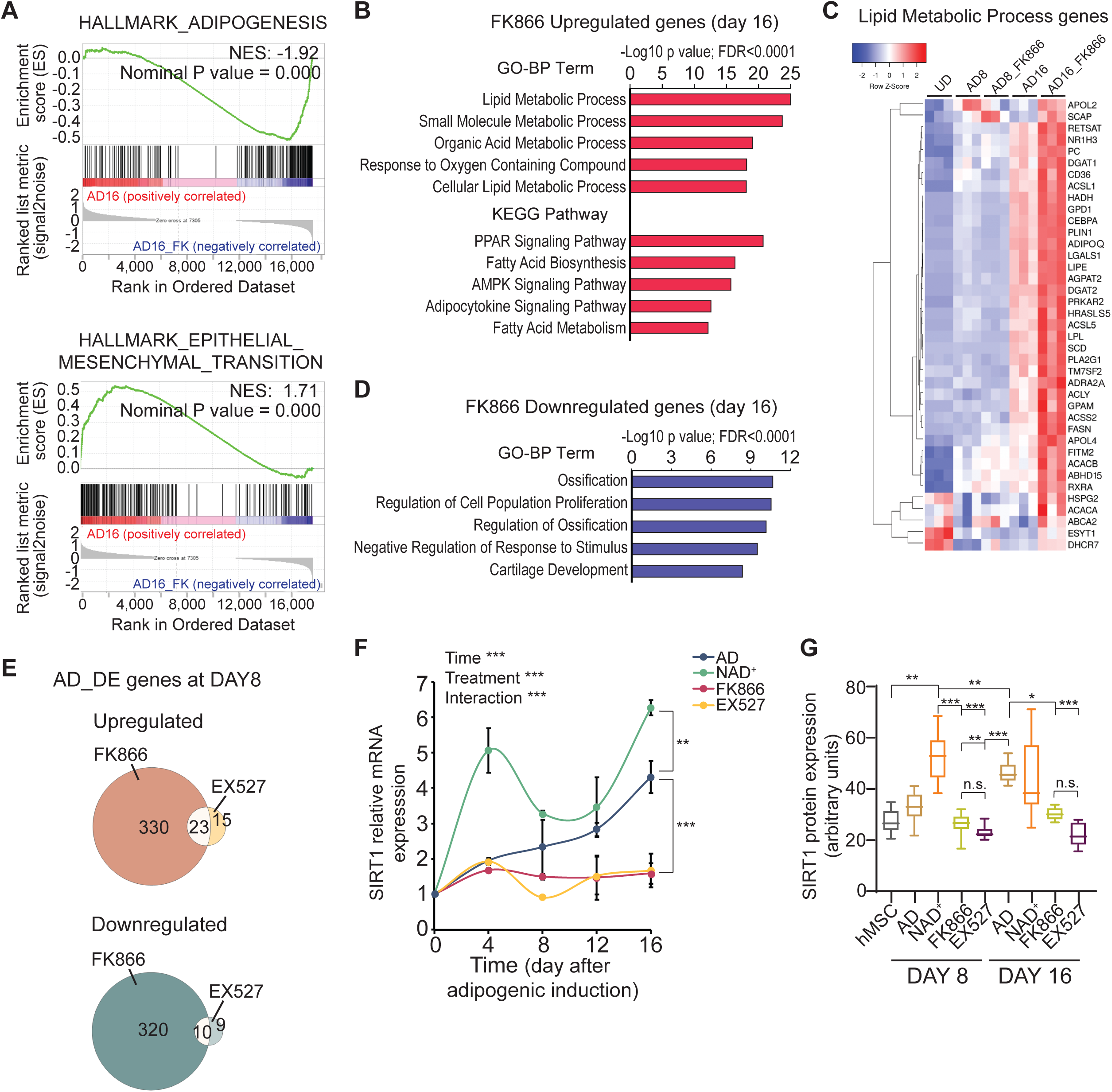
The NAD^+^ salvage pathway is dispensable for adipogenesis. **A)** Gene set enrichment analysis (GSEA) was investigated within the molecular signature database (MSigDB) “Hallmark” gene set collection. Genes were rank-ordered by differential expression between terminally differentiated adipocytes untreated (AD16) or treated with 1 nM FK866 (AD16_FK). **B)** Functional annotation (Biological processes and KEGG pathways) for 144 genes upregulated by FK866 treatment during adipogenesis, at terminal adipogenic differentiation (day 16). **C)** Heatmap comparing expression levels between the indicated samples at day 16 form 39 genes involved in lipid metabolism. **D)** Biological processes enriched in 44 genes downregulated by FK866 treatment during adipogenesis, at terminal differentiation (day 16). **E)** Overlap of DE genes (up- or downregulated) comparing EX527 and FK866 treated cells during adipogenic induction, at day 8 **F)** *SIRT1* gene expression levels were assessed by RT-qPCR at the indicated days after adipogenic induction on hMSC either untreated (AD) or treated with the indicated drugs. n= 3 biological and 2 technical replicates. One-way ANOVA followed by Tukeýs post test. ** p <0.01, *** p <0.001. **G)** Boxplot showing SIRT1 protein levels analyzed by immunofluorescence at days 8 and 16 after adipogenic induction on hMSC. Cells were either untreated (AD), or treated with the indicated compounds. Densitometric analyses are from n= 2 biological and 7 technical replicates. Kluskal-Wallis test followed by Dunn’s multiple comparisons test was applied. n.s.: non-significant * p <0.05, ** p <0.01, *** p <0.001.

### Increased NAD^+^ bioavailability during the adipogenic process impairs the rise of mitochondrial respiration capacity in hMSCs

A major shift in metabolic phenotype is a hallmark of adipogenic differentiation. Hence, we investigated the functional effect of altering NAD^+^ balance and SIRT1 activity during hMSC adipogenesis on energy metabolism by performing extracellular flux analysis for measuring cellular bioenergetics. As expected, non-treated hMSCs progressively increased their respiratory capacity during the adipogenic differentiation when compared to undifferentiated hMSCs, which retained low oxygen consumption rates across all tested days (Figure 5A-D). We observed that at day 4, all tested conditions retained low respiratory capacity, comparable to undifferentiated cells (Figure 5A), while the most prominent increase in respiration capacity occurs between days 8 to 16 (Figure 5B-D). Indeed, FK866 treatment overall allowed the metabolic reprogramming during adipogenesis of hMSCs; however, NAD^+^ treatment obstructed the progressive increase in mitochondrial respiration (Figure 5A-D). Interestingly, pharmacological inhibition of SIRT1 with EX527 showed major effect after day 12, consisting of markedly reduced respiratory capacity compared to the untreated cells. These results confirm that metabolic reprogramming during adipogenesis is compromised by SIRT1 inhibition specifically at late differentiation stages and reinforce the notion that SIRT1 is essential for adipocyte maturation. We observed major differences between treatments in maximal respiration and spare respiratory capacity at days 12 and 16, when induced cells either untreated or treated with FK866 showed a very significant increase compared to the rest of the conditions (Figure 5E-H, P<0.0001, Two-way ANOVA with Tukey’s post-test), indicating a high rate of oxidative phosphorylation in these cells. Non-mitochondrial respiration and proton leak did not show significant differences at any of the studied conditions (Figure S5). Notably, extracellular acidification rate (ECAR) measurements revealed that undifferentiated hMSC exhibit a glycolytic phenotype, while treatment with NAD^+^ during adipogenic induction also diminished the glycolytic flux (Figure 5I-L), indicating that these cells are metabolically less active.

**Figure 5:**
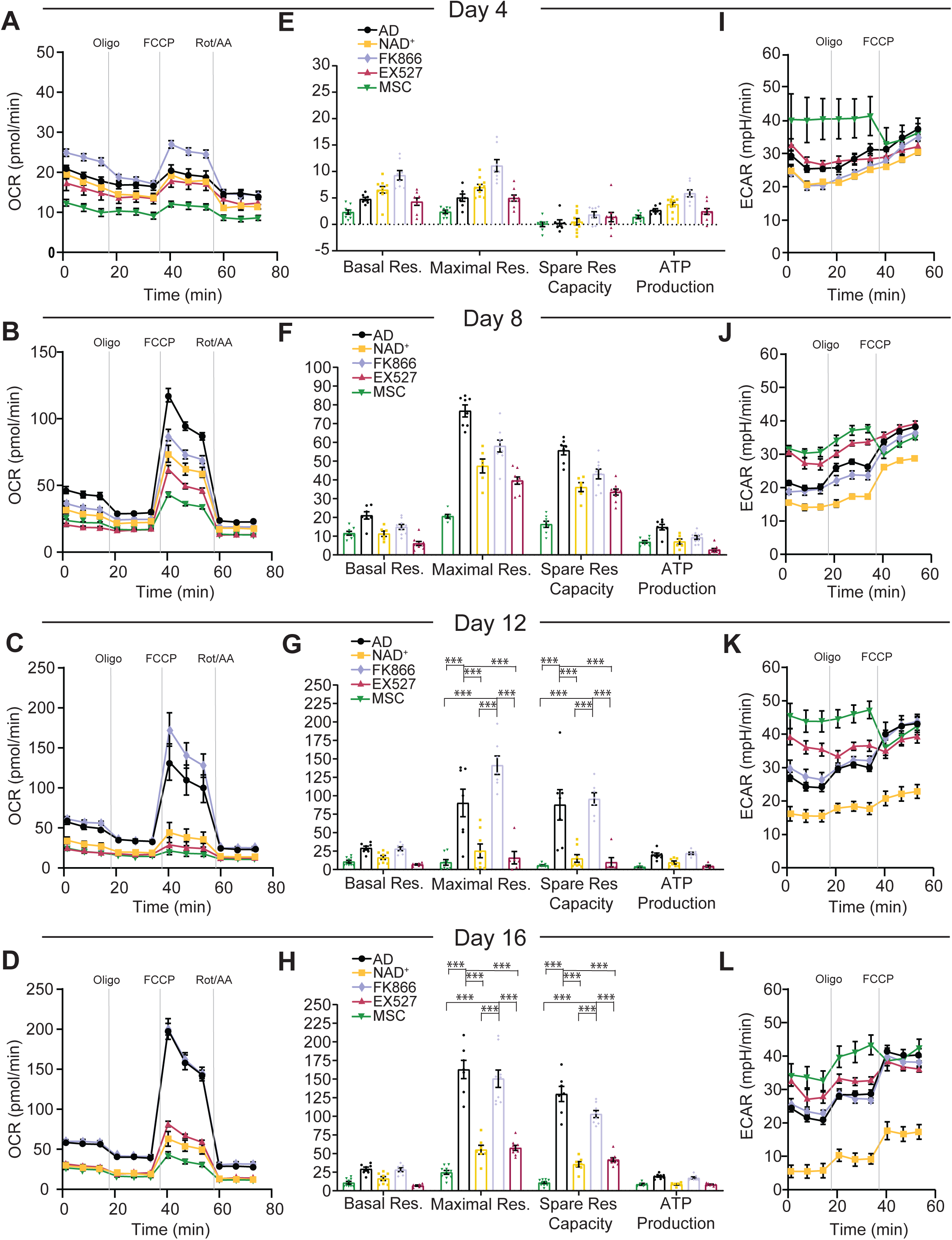
NAD+ impairs mitochondrial bioenergetics during adipogenic induction in hMSC. Analysis of oxygen consumption rate (OCR) and extracellular acidification rate (ECAR) was performed using Seahorse XF analyzer to assess mitochondrial respiration and lactate production from n = 3 biological replicates with 6-10 technical replicates each. **A-D)** OCR was measured at days 4 (A), 8 (B), 12 (C) or 16 (D) after adipogenic induction in hMSC, in the absence or presence of the indicated treatments. with sequential addition of oligomycin (Oligo, complex V inhibitor), FCCP (a protonophore), and Rotenone/antimycin A (Rot/AA, complex III inhibitor), **E-F**) Mitochondrial bioenergetic parameters calculated from extracellular flux analyses: basal respiration, maximal respiratory capacity, spare respiratory capacity, and ATP production. Two-way ANOVA followed by Tukeýs post test. * p <0.05, ** p <0.01, *** p <0.001. **I-L)** ECAR was measured after serial addition of oligomycin and FCCP. Data is presented by mean ±SEM. AD: adipogenic induced cells; NAD+ adipogenic induced cells treated with 5 mM NAD^+^; FK866: adipogenic induced cells treated with 1 nM FK866; EX527: adipogenic induced cells treated with 50 μM EX527. MSC: untreated, undifferentiated hMSC.

Next, we performed label-free two-photon fluorescence lifetime microscopy (2P-FLIM) of the intrinsic metabolic biomarker NADH in live cells (see Methods section). We examined the fraction of bound NADH (fB_NADH) during adipogenic differentiation with all treatments with a micrometer pixel resolution (Figure 6A). The results were in agreement with the extracellular flux analyses, and we saw that the fB_NADH progressively increased during adipogenic differentiation, as a result of the metabolic shift from a glycolytic to an OXPHOS phenotype (Figure 6A, S6A-E). Also, NAD^+^ treatment induced consistently low fB_NADH across all tested days (Figure 6A), reinforcing the notion that increased NAD^+^ bioavailability during adipogenic differentiation opposes the metabolic shift towards OXPHOS. With this approach, we only found significant reduction of fB_NADH in cells treated with EX572 at the end of the differentiation process (Figure 6A), which is in line with the extracellular flux analyses indicating that SIRT1 inhibition hinders adipocytic maturation.

**Figure 6:**
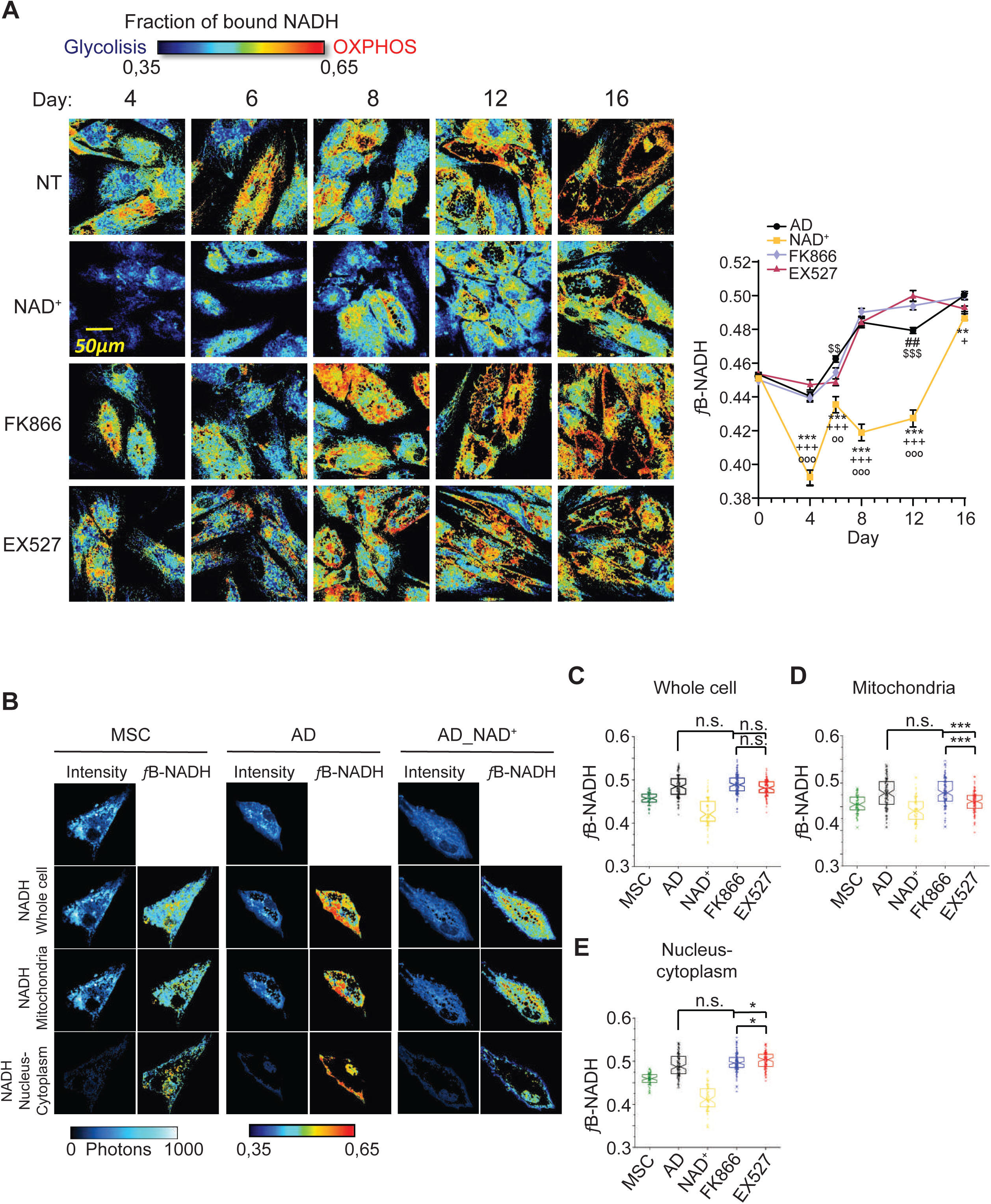
Subcellular compartmentalization of NADH metabolism during adipogenesis depends on SIRT1 activity. **A)** Representative images of fraction of bound NADH (fB_NADH) of hMSCs during adipogenic differentiation at days 4, 6, 8, 12 and 16 of induction in the absence (NT) or presence of the indicated treatments: 5 mM NAD^+^, 1 nM FK866 or 50 μM EX527. Low fB_NADH (blue colors) corresponds to a cellular glycolytic phenotype, while high fB_NADH (red colors) corresponds to an OXPHOS phenotype. Quantification of fraction of bound NADH in each culture condition was performed from n= 14 single cells. Experiments were conducted per triplicate. Data is presented by mean ±SEM. **B)** Representative images of intensity and fB_NADH of hMSCs, pre-adipocytes (AD) and cells treated with NAD+ during adipogenic induction (AD_NAD^+^) imaged at day 8 of differentiation, show different spatial distributions of fraction of bound NADH in different cell compartments such as mitochondria, nucleus and cytoplasm. **C-E)** Quantification of fB_NADH in single cells at day 8 from hMSC (MSC), pre-adipcyte (AD) and cells treated with the indicated compounds during adipogenic differentiation. Quantification from n= 63-125 cells was performed in the whole cell (C), or the mitochondrial (D) and nuclear/cytoplasmic (E) subcellular compartment. Two-way ANOVA followed by Tukeýs post test. * p <0.05, ** p <0.01, *** p <0.001, n.s.: non-significant.

### Subcellular compartmentalization of NADH metabolism during adipogenesis depends on SIRT1 and is impaired by abnormally high NAD^+^ levels

Our FLIM and extracellular flux analyses showed important disparities specifically for the cells treated with EX527 during adipogenic differentiation. Interestingly, at days 8 and 12, two photon-FLIM didńt show significant differences between EX527 and untreated cells, while extracellular flux analyses indicated an important reduction in respiratory capacity of EX527-treated cells after day 8. For this reason, we ought to determine the metabolic signatures at distinct subcellular compartments, in order to dissect^i^ the subcellular location of the metabolic changes in these cells (see Methods section and Figure S1). This technique allowed us to capture intensity images and maps of fB_ NADH in the entire cell and in different cellular compartments such as mitochondria and nucleus/cytoplasm, as shown in Figure 6B for a hMSC, and at day 8 after adipogenic induction in untreated (AD) or NAD^+^ treated cells (AD_NAD^+^). Our single cell analysis at day 8 showed a wide distribution of cellular metabolic states in every culture condition (UD, AD, AD-NAD+, AD-FK866 or AD-EX527) (Figure 6D). Considering the metabolic fingerprint of the entire cell, NAD^+^ treated cells showed lower fB_NADH than untreated cultures, while FK866 and EX527 treatments didńt show differences with untreated cells (Figure 6C, Two-way ANOVA with Tukey’s post-test). Interestingly, we observed distinct subcellular compartmentalization of NADH metabolism depending on the treatment (Figure 6D, E). Consistently, fB_NADH at both mitochondria and nucleus/cytoplasm were lower in NAD^+^ treated cells (Figure 6D-E). Moreover, the fB_NADH in the cytoplasm/nucleus was higher than in mitochondria in differentiating cells, while remained similar in hMSC; yet NAD^+^ treatment showed an opposite ratio, where the mitochondrial fraction showed significantly higher levels of fB_NADH (Figure S6F). Interestingly, the fB_NADH of mitochondria appeared significantly decrease in cells treated with the SIRT1 inhibitor, while the fB_NADH increased within the nucleus/cytoplasm compartment in these cells when compared with the non-treated or with the FK866 treated cells (Figure 6D,E; P<0,05, Two-way ANOVA with Tukey’s post-test). Together, out data reveal that energy metabolism progress during adipogenic differentiation is impeded by both SIRT1 inhibition and high NAD^+^ levels, through different mechanisms at distinct subcellular compartments. These observations have further implications for disease, as provide previously unappreciated sub-cellular insights into the previously reported efficacy of NAD^+^ as a treatment for diet-induced obesity and metabolic dysfunction(39, 40).

## DISCUSSION

In this study, we have demonstrated a previously unappreciated role for NAD+-SIRT1 interplay in adipogenesis from hMSC which is dependent on the differentiation stage. We have shown that SIRT1 activity is essential for terminal adipocyte differentiation, and unexpectedly, NAD^+^ availability fine tunes the adipogenic process. Mounting research demonstrates that SIRT1 inhibits adipogenesis in mesenchymal stem cells(41–47), however the dynamics of SIRT1 activity during adipogenesis remains poorly understood. Our gene expression data reveals that SIRT1 activity is dispensable for adipocyte commitment, as out of ∼5,000 DE genes at day 8 of differentiation, less than 60 were significantly altered between normally differentiating cells and those with inhibited SIRT1 activity. Unexpectedly, transition to mature adipocytes was strongly dampened by SIRT1 inhibition, demonstrating SIRT1 is essential. These results are in line with the notion that reducing SIRT1 activity specifically in fat could lead to improved metabolic function in obesity(48) . Moreover, in mice lacking SIRT1 specifically in MSC, their adipogenic capacity appears compromised leading to significant reduction in subcutaneous fat(49), further reinforcing this notion.

A number of studies report that the stem cell redox status is tightly regulated during differentiation, and thus activation of oxidative pathways constitute a metabolic signature of stem-cell differentiation(50, 51). Along the same lines, stem cells appear to contain lower levels of reactive oxygen species (ROS) than their mature progeny, and that ROS accumulation triggers intracellular signaling required for differentiation(52–55). Hence, it appears that metabolic reprogramming activates specific signaling cascades promoting either stem cell maintenance and self-renewal (reduced state) or stem cell proliferation and differentiation (oxidized state). Here, we have shown that fueling energy metabolism with NAD^+^ markedly obstructs adipogenic differentiation. Imposing an oxidative redox state during early differentiation triggers a transcriptional program leading to translational arrest and induction of proapoptotic pathways, and these cells acquire a quiescent metabolic phenotype (Figure 2, 4). NAD^+^ levels rise at late stages of differentiation(23), probably as a result of the increased oxidative metabolism which in our hMSC adipogenic differentiation model rapidly emerges between days 8-12 (Figure 5, 6A). This is coincident with increased levels of SIRT1 transcription and protein expression (Figure 4F, 4G), reinforcing the idea of a SIRT1-dependent late stage of adipogenic differentiation. At this regard, obstructing the NAD^+^ salvage pathway through constant inhibition of the rate-limiting enzyme NAMPT during adipogenesis didńt hinder the adipogenic capacity, demonstrating that the metabolic switch triggered by the transcriptional rewiring might control intracellular variations on NAD^+^ levels and activity of NAD^+^- consuming enzymes. Indeed, we observed that NAMPT inhibition upregulated transcription of key lipid metabolism genes, and adipocytic identity at terminal differentiation (Figure 4A-C), thus favoring lipid accumulation (Figure 1A). This suggests that the NAD^+^ salvage pathway fine tunes adipogenesis in its late stages, and is in line with its protective role in diet-induced obesity(56, 57).

To gain insights into our observations in live, single cells at a submicrometric resolution, we developed a new method based on two-photon fluorescence lifetime microscopy (2P-FLIM) of intrinsic fluorophores that provides directly quantitative metrics of adipogenic stem cell differentiation, metabolic state of subcellular compartments. We measured a long lifetime of lipid droplets associated fluorophores (Figure 1B, 1C), as previously shown(58). We used the lifetime contrast to select autofluorescence from lipids and from NADH within cells (Figure S1A, S1B). Third harmonic generation microscopy was performed to visualize lipid droplets (26, 59) . Based on NADH intensity contrast, we also implemented an automated segmentation of mitochondria and cytoplasm/nuclear compartments that relies on their different concentration of NADH (Figure S1B). With this method, we observed lipid accumulation and increased fraction of bound NADH during adipogenic differentiation in live single cells, reflecting the metabolic shift from a glycolysis to OXPHOS metabolism during differentiation(60, 61). We provided here the first description of intracellular NADH metabolic signature in different subcellular compartments and at distinct stages of adipogenic differentiation in human cells, since subcellular characterisation of NAD^+^/NADH metabolism was previously demonstrated during few hours after adipogenic induction from 3T3-L1 murine preadipocytes(62, 63). Our results show that spatial subcellular compartmentalisation of NADH metabolism is highly dependent on the differentiation stage and is intimately linked with the transcriptional reprograming allowing progressive lipid accumulation. Hereby, our observations are in agreement with the emerging view that temporal and spatial subcellular metabolic compartmentalisation contributes to numerous biological roles and regulation of intracellular signalling and transcription (24, 64) and that NAD^+^ compartmentalisation regulates adipogenesis(63).

Finally, the quantitative metrics based on fluorescence lifetime microscopy developed in this study serves as a label-free biomarker to simultaneously measure lipogenesis and metabolic shifts in single cell. Quantitative characterization of subcellular states from adipose tissues in health and disease using our two-photon microscopy-based method could provide means to uncover new roles of hMSC in obesity, thus paving the way for the development of MSC-based treatments.

## ACKNOWLEDGEMENTS

We thank all members of Aguilar-Arnal and Chiara Stringari laboratories for helpful discussions and advice. We thank Pierre Mahou, Isabelle Lamarre and Emmanuel Beaurepaire for scientific discussions and technical help. We are thankful to Marcia Bustamante Zepeda, MSc, and Miguel Tapia Rodríguez, at the IIBo, for technical assistance. We are thankful to Dr. Alfonso Leon-del-Rio laboratory at the IIBo and particularly to Salvador Ramírez Jiménez, MSc, for kindly sharing reagents and equipment. This work was supported by Human Frontier Science Program (HFSP) under the contract RGY0078/2017 ChroMet to CS and LA-A, the Agence Nationale de la Recherche (ANR) under contracts ANR-10-INBS-04 France BioImaging and ANR-11-EQPX-0029 Morphoscope2 to CS, and grant PAPIIT IN210619 from Universidad Nacional Autónoma de México (UNAM) to LA-A. RO-S lab was supported by National Council of Science and Technology (CONACyT) (grants FC 2016/2672 and FOSISS 272757). ES-R acknowledges the reception of PhD fellowship from CONACyT and the PhD program Doctorado en Ciencias Bioquímicas-UNAM.

## AUTHOR’S CONTRIBUTION

LA-A and CS conceived and designed the study. ES-R, TPLU, XdT-R, GF-O conducted experiments. ES-R, TPLU, XdT-R, RO-S, CS and LA-A analyzed and interpreted the data. LN and AT assisted with Seahorse extracellular flux analyses. JJM provided hMSC and validated the model. Two-photon fluorescence lifetime microscopy was performed in CS lab and molecular biology and RNA-seq were done in LA-A lab. LA-A, CS, TPLU, ES-R and GF-O wrote the manuscript. All authors reviewed the manuscript.

## DECLARATION OF INTEREST

The authors declare that the research was conducted in the absence of any commercial or financial relationships that could be construed as a potential conflict of interest.

## DATA AVAILABILITY

All relevant data to this manuscript are available from the authors.

## METHODS

### Isolation and characterization of BM-MSCs

Bone marrow derived MSCs were obtained from healthy donors and have been previously described(65). Briefly, mononuclear cells (MNCs) from BM were obtained by density gradient centrifugation, and 2×10^5^ MNCs/cm^2^ were seeded in low glucose Dulbecco’s modified Eagle’s medium (LG-DMEM, Gibco) with 10% fetal bovine serum (FBS; Gibco), 4mM L-glutamine and antibiotics. Cells were incubated at 37°C and 5% of CO_2_. At 80% of confluence, adherent cells were trypsinized and reseeded at a density of 0.2×10^4^ cells/cm^2^.

Experiments were done at 3 - 5 passages. Cell surface markers expression of MSCs were determined by flow cytometry following our previously implemented and described protocols (65, 66), where MSCs were selected to express CD73, CD90 and CD105 markers, while being negative to the hematopoietic markers CD45, CD34 and CD14 (Figure S7A). Differentiation capacity was assessed using the StemPro™ Adipogenic and Osteogenic differentiation kits (Gibco A1007001, A1007201), and the Chondrogenic Differentiation Medium (Cambrex Bio Science) supplemented with 10 ng/ml of TGFβ (Peprotech 100-21C), following manufactureŕs instructions. Adipogenic differentiation was evaluated by Oil Red O (Sigma O0625) staining after 16 days of induction, osteogenic differentiation was revealed by detecting alkaline phosphatase activity (Sigma B5655) after 14 days of induction, and chondrogenic differentiation was indicated by the presence of mucopolysaccharides positive to alcian blue (Sigma, ca. no. A5268), in micromasses cross-sections, after 28 days of induction(65, 66). Representative images are shown in Figure S7B.

All protocols were compliant with the Declaration of Helsinki and approved by the Ethics Committee of Villacoapa Hospital, Mexican Institute for Social Security (IMSS). Informed consent was given by the participants. Additionally, a BM-MSC line was purchased from ATCC .

### Cell culture and maintenance

hMSCs were maintained in LG-DMEM (Gibco, cat. no. 31600-034) supplemented with 10% FBS, 4mM L-glutamine, 100 μg/ml of penicillin and streptomycin (Gibco), and incubated at 37°C and 5% of CO_2_. Adipogenic differentiation was induced in MSCs in growing phase (70%-80% confluency), and drugs were present in the medium as indicated in the text and figures, following our previously standardized protocols(24). The medium and drugs were replaced every four days during the differentiation process. For live cell imaging, BM-hMSCs (ATCC) were cultured on 3.5- cm glass bottom petri dishes (MatTek, Ashland, MA, USA) in 2.5 ml LG-DMEM per well without phenol red and 10% FBS and 100 UI/mL penicillin and 100 µg/ml streptomycin. Cells were plated at the initial density of 1,1×10^4^ cells/cm^2^ and allowed to attach overnight in a humidified cell culture incubator at 37 °C in 5% CO_2_ before proceeding with treatments. During the imaging experiments, we replaced the adipogenic medium by basal medium without the phenol red before imaging. To determine the metabolic trajectory of NADH lifetime, MSCs were treated with rotenone 50 µM in DMSO (R8875; Sigma-Aldrich, St. Louis, MO, USA) and hydrogen peroxide (H_2_O_2_) 4mM in DMSO (216763; Sigma-Aldrich, St. Louis, MO, USA) to block the respiratory chain via complex I and to increase NAD^+^:NADH ratio via oxidative stress respectively. All cultures tested negative for mycoplasma contamination.

### Antibodies and reagents

The antibodies used in this study are as follows: anti-SIRT1, Millipore cat. no. 07-131; anti-PPARγ, Cell Signaling, cat. no. 81B8; anti-GAPDH-HRP conjugated, GeneTex, cat. no. GTX627408-01; Goat anti-Rabbit Alexa Fluor® 594 conjugated, Invitrogen cat. no. R37117; anti rabbit IgG-HRP conjugated, Invitrogen cat. no. 65-6120. All purchased antibodies were validated for mammalian studies (as shown on the manufacturers’ websites). EX527 (E7034), FK866 (F8557) and NAD^+^ (N8535) were purchased from Sigma.

### Oil-Red-O staining

Cells grown on slides were briefly washed with PBS and fixed for 45 min with 4% fresh paraformaldehyde. Preparation of Oil Red O (SIGMA, cat. no.O1391) working solution and staining of slides were performed as described(67). ORO was applied on the slides for 5 min at RT. Slides were washed twice during 10 min. in water, and mounted in vectashield mounting media (Vector Labs, cat. no. H-1000). The images were captured with the camera Axiocam EEc 5s coupled to a ZEISS Primovert microscope, using a 20X magnification. Lipid droplets were quantified using the ImageJ software, by converting RGB to 8-bit grayscale images, and then using the “analyze particles” plug-in as described(68). Four frames per slide were used for image analyses and quantification (n=3 biological replicates with 4 technical replicates).

### Quantitative real-time PCR

Total RNA was extracted from MSCs using TRIzol™ Reagent (Invitrogen cat. no. 15596018) following the manufactureŕs instructions. cDNA was obtained by retrotranscription of 1 μg of total mRNA with iScript cDNA synthesis kit (Bio-Rad) according to the manufacturer’s instructions. Real-time RT-PCR was done with the real-time CFX96 detection system (Bio-Rad). For a 10 μl PCR reaction, 25 ng of cDNA template was mixed with the primers to final concentrations of 200 nM and mixed with 5 μl of iTaqTM Universal SYBR® Green Supermix kit (Bio-Rad). The reactions were done in triplicates with the following conditions: 30 sec at 95 °C, followed by 45 cycles of 30 s at 95 °C and 30 s at 60 °C. Expression levels were calculated using the ddCt method. The PCR primers were as follows: SIRT1: Fw, 5′-GCTGGAACAGGTTGCGGGAA-3′; Rv, 5′-GGGCACCTAGGACATCGAGGA-3′. β-actin: Fw, 5′-CTTGTACGCCAACACAGTGC-3′; Rv, 5′-ATACTCCTGCTTGCTGATCC-3′

### Immunofluorescences

MSCs were seeded on Lab-Tek Chamber slides (Thermo Fisher) at 9×10^3^ cells/cm^2^. Cells were washed with phosphate buffered saline (PBS; Gibco), fixed with 1% of paraformaldehyde at 37°C for 10 min, washed twice with PBS and permeabilized with PBS 0.1% triton during 15 min. The slides were then washed with PBS, and incubated with blocking buffer (PBS, 0.1% Tween® 20, 2% BSA) for 1 hour. Incubation with anti-SIRT1 (1:500) was performed over night at 4°C, and the secondary antibody (1:2,000) was incubated during 1 hour at RT. 1:5,000 dilution of Hoechst 33342 was used for nuclear counterstain (Invitrogen H1399) by incubating during 10 min at RT. Coverslips were mounted using VECTASHIELD® antifade mounting medium (Vector labs H1000) and sealed with nail polish. Fluorescence images were acquired by an Olympus DP70 Digital Camera in an Olympus BX51 fluorescence microscope. Hoescht stain was acquired at a 1/300 s exposure, while SIRT1 intensity was acquired at 1/200 s. Densitometry was performed using ImageJ on 3 cells at 4 different fields from 2 biological replicates.

### Western blotting

MSCs were harvested from confluent 6-well dishes, washed with PBS and lysed with RIPA buffer supplemented with HDAC inhibitors (50mM Tris pH 8, 150mM NaCl, 1% 5mM EDTA pH8, 1% NP-40, 0.5% Na deoxycholate, and 0.1% SDS; all from Sigma). Cells where left on ice during 20 min. and centrifuged at 14,000 RPM during 15 min at 4°C. The protein extracts in the supernatants were snap frozen and stored at -80°C. Protein quantification was done using the Bradford colorimetric assay (Sigma B6916). 20µg of proteins were separated run on a 10% SDS-PAGE gel, at 100 V for 2 hours using a mini-PROTEAN system (BIO-RAD) and transferred to nitrocellulose membrane (Millipore) at 40 mV overnight at 4°C. A 1:1,500 dilution of anti-PPARγ in blocking buffer (Tris-buffered saline plus Tween-20 -TBST- and 5% nonfat milk) was used to detect the protein. Antibodies to GAPDH protein at 1:20,000 dilution in PBST were used as a loading control. Proteins were revealed using chemiluminescent detection (Immobilon Western, Millipore WBKLS0100) and visualized using a Kodak GEL Logic 1500 Imaging System with Transilluminator.

### Extracellular flux analysis

OCR and ECAR were measured using the Seahorse XFe96 Analyzer (Agilent), using The Cell Mito Stress Test Kit (Agilent, cat. no. 103015-100). MSCs were seeded at a density of 10,000 cells/well in a XF96 cell culture 96-well microplate (Agilent 101085-004) precoated with 10 µg/ml of fibronectin (Sigma F1141). Adipogenic induction and treatments were initiated after two days of seeding. One hour prior the Seahorse analysis, MSCs culture were washed with 200 µl/well of XF assay media supplemented with 10 mM glucose, 2mM glutamine and 1 mM pyruvate. Then, 180 µl/well of this medium was added and the plate was equilibrated for 30 min at 37°C in a CO_2_-free incubator before being transferred to the Seahorse XFe96 analyzer. Measurement of OCR and ECAR was done at baseline and following sequential injections of (i) 2 μM oligomycin, an ATP synthase inhibitor, (ii) 0.5 μM carbonyl cyanide-4-(trifluoromethoxy) phenylhydrazone (FCCP), a protonophoric uncoupler, and (iii) 0,5 μM of rotenone, an inhibitor of complex I of the electron transport chain. Briefly, oligomycin inhibits mitochondrial ATP synthase, and the resulting drop in OCR and rise in ECAR are attributed to ATP-linked OCR and the compensation of glycolysis for the loss of mitochondrial ATP production. FCCP uncouples the mitochondrial proton gradient and oxygen consumption from ATP synthase, hereby driving maximal OCR. Rotenone inhibits complex I of the electron transport chain, hence it hinders mitochondrial oxygen consumption. Therefore, the residual OCR is regarded as nonmitochondrial.

### Expression profiling (RNA-seq) and analysis

Total RNA from MSCs was extracted using Quick RNA MiniPrep Kit (Zymo Research, USA) following the manufactureŕs instructions. RNA samples with RNA integrity number (RINs) > 7.0 were sent for library preparation and sequencing to Novogene Corporation Inc., California, USA. Briefly, mRNA was isolated using oligo(dT) beads and randomly fragmented by adding fragmentation buffer, followed by cDNA synthesis primed with random hexamers. Next, a second-strand synthesis buffer (Illumina), dNTPs, RNase H and DNA polymerase I were added for second-strand synthesis. After end repair, barcode ligation and sequencing adaptor ligation, the double-stranded cDNA library was completed with size selection to 250-300 bp, and PCR enrichment. Sequencing was performed on an Illumina NovaSeq 6000 Sequencing System with paired-end 150 bp reads, at 9 G raw data/sample. Total and mapped reads per sample are shown in Supplementary Table S1.

### RNA-seq data processing

Human Reference genome and gene model annotation files were downloaded from genome website browser (NCBI/UCSC/Ensembl) directly. Indexes of the reference genome was built using STAR and paired-end clean reads were aligned to the *Homo sapiens* assembly GRCh38/hg38, with the STAR aligner v2.5 (69). STAR uses the method of Maximal Mappable Prefix (MMP) which can generate a precise mapping result for junction reads. HTSeq v0.6.1 was used to count the read numbers mapped of each gene. Afterwards, FPKM of each gene was calculated based on the length of the gene and reads count mapped to it (70). Differential expression analysis between conditions (three biological replicates per condition) was performed with the DESeq2 R package (2_1.6.3), which uses a model based on the negative binomial distribution (32). The resulting *P*-values were adjusted using the Benjamini and Hochberg’s approach for controlling the False Discovery Rate (FDR). Genes with an adjusted *P*-value <0.05 were assigned as differentially expressed. The Venn diagrams were prepared using the function vennDiagram in R based on the gene list for different groups, or with Venny V 2.1 (https://bioinfogp.cnb.csic.es/tools/venny/). Differentially expressed genes were subjected to functional analyses using the “Compute Overlaps” tool to explore overlap with the CP (Canonical Pathways) and the GO:BP (GO biological process) gene sets at the MSigDB (molecular signature database). The tool is available at: https://www.gsea-msigdb.org/gsea/msigdb/annotate.jsp, and estimates statistical significance by calculating the FDR q-value. This is the FDR analog of the hypergeometric P-value after correction for multiple hypothesis testing according to Benjamini and Hochberg. Gene set enrichment analysis (GSEA) was performed using GSEA v. 4.0.3. (71) to determine the enrichment score within the Hallmark gene set collection in MSigDB v7.0(72), selecting the Signal2Noise as the metric for ranking genes. The findMotifs.pl program in the HOMER software (^73^) was used for motif discovery and enrichment, searching within the genomic regions encompassing 300 Kb upstream and 50 Kb downstream the TSS, and selecting 6-8 bp for motif length. Motif enrichment is calculated by findMotifs.pl using the cumulative hypergeometric distribution.

All RNA-seq raw and processed data will be made publicly available at Gene Expression Omnibus (GEO) under the accession number GSE178615

### Two-photon excited fluorescence lifetime imaging (FLIM) and third harmonic generation (THG)

Imaging was performed on a laser scanning microscope (TriMScope, Lavision Biotec, Germany). A simplified scheme of the multiphoton microscope is shown in Figure S7C. The Excitation is provided by a dual-output femtosecond laser (Insight DS++, Spectra-Physics, Santa Clara, CA, USA) with a first beam turnable from 680 nm to 1300 nm (120 fs pulses, 80 MHz) and a second, fixed wavelength beam at 1040 nm (200 fs pulses). A water Immersion objective (25X, NA=1.05, XLPLN-MP, Olympus, Japan) is used to focus the laser on the sample and collect fluoresce signal. Fluorescence signal is epi-detected by a hybrid photomultiplier tube (R10467U, Hamamatsu, Japan -), whereas and third-harmonic generation (THG) signal is forward detected by a photomultiplier (H6780-01, Hamamatsu, Japan). To perform Fluorescence lifetime microscopy of NADH, 760 nm wavelength excitation was used with a typical power of 12 mW (Figure S7D). A band-pass filter was used in front of the detector to collect NADH autofluorescence (Semrock FF01–460/80).Time-correlated single photon counting (TCSPC) electronics (Lavision Biotec, Germany) with 5,5 ns dead time, and 27 ps time bins was used to measure the arrival time of the fluorescence photons with respect to the laser pulse and perform FLIM imaging. The laser trigger reference was taken from the fixed wavelength beam using a photodiode (PDA10CF-EC, Thorlab). Calibration of the FLIM system was performed by measuring the lifetime of fluorescein at pH=9 with a single exponential of 4 ns (Figure S7E). We measured the lifetime of free NADH in solution (Sigma Aldrich n. N8129, St. Louis, MO, USA) to calculate the fraction of bound NADH (Figure S6B). We typically collected 500 photons for FLIM images of live cells with a pixel dwell time of 240 μs/pixel and a total acquisition time on the order of one minute. Third harmonic generation was performed using a wavelength of 1100nm with a typical power of 12mW and the signal was collected with a band-pass filter (Semrock FF01–377/50) (Figure S7D). We typically collected 800 photons for THG images with a a pixel dwell time of 53 μs/pixel and a total acquisition time of the order of one minute.

### Analysis of the Fluorescence Lifetime microscopy images

Intensity images were analysed with Fiji-ImageJ (NIH, Bethesda, MD, USA). All FLIM data was processed and analysed with SimFCS (developed by the laboratory for Fluorescence Dynamics https://www.lfd.uci.edu/globals/) and with a Matlab (Mathworks, Natick, MA, USA) custom written software. FLIM data were transformed by using FFT and plotted in the phasor plot as previously described (74) (75) (see Supplementary material). Briefly, the coordinates *g* and *s* in the phasor plot were calculated from the fluorescence intensity decay of each pixel of the image (Figure S7F) by using the transformations defined in the Supplementary material (equation 1 and 2). We applied an intensity threshold to eliminate the background of the cellular medium and a median filter on the *g* and *s* images to reduce the variance of the phasor location without decreasing the spatial resolution of the image. For every pixel of the image, we calculated the value of *τ _φ_* (equation (5) in Supplementary material) and *τ _m_* (equation (6) in Supplementary material) starting from the g and s images (Figure S1A). Fraction of bound NADH (equation (7) in Supplementary material) was graphically calculated as the distance from the experimental point to the location of free NADH (Figure S1A and Figure S6B).

### Subcellular segmentation of lipid droplets, cell NADH, mitochondria and nucleus and cytoplasm

Image processing and segmentation was performed by a Matlab custom written software. The principles of the segmentation are illustrated in Figure S1B. A threshold (2.87 ns) was applied in the τ_m_ lifetime image to automatically separate lipid droplets and NADH of the cell. Pixels with longer lifetime were assigned to a lipid mask while pixels with shorter lifetime were highlighted to the NADH cell mask. The τ_m_ threshold was determined empirically to match the lipid mask border with the THG signal of the lipid droplet (Figure 1B). Then the mask of NADH cells was used to calculate fB_NADH of the same ROI or cell. For quantification, we used the average values of fB_NADH and lipid ratio. To quantify the lipid droplets in ROIs or in single cells, we calculated the ratio between the number of pixels of the lipid mask and the total number pixels of the cell: Single cell analysis was performed manually. We performed subcellular segmentation of mitochondria and nucleus plus cytoplasm applying a threshold (150 photons) to the intensity image multiplied with the NADH mask (Figure S1B). The threshold was determined based on the different NAD^+^/NADH ratios in mitochondria (∼ 10) and cytoplasm and nucleus (∼ 50-1000)(76). Pixels with higher number of photons were assigned to mitochondria while pixels with lower number of photons are assigned to nucleus and cytoplasm. The masks of mitochondria and nucleus and cytoplasm were applied to the map of fraction of bound NADH, and the average fraction of bound NADH was calculated in different cellular compartments for statistical analysis.

### Statistics and images analyses

Data are shown as mean with SEM. Statistical analyses were performed using GraphPad Prism 8.2. The statistical tests were performed as indicated in the figure legends, mostly consisting of Two way ANOVA followed by Turkeýs multiple comparisons test, or Kruskal–Wallis H test . Statistical significance was considered when the *P* value was <0.05. When possible, experimental evaluation was performed blind to the experimental conditions (i.e., specifically for western blot quantification, image processing and subsequent quantifications).Western blot analyses and image processing for overlay in different channels from immunofluorescences were performed with ImageJ software. Figures were arranged using Adobe Illustrator.

## Supplementary Information

**Fig. S1.**
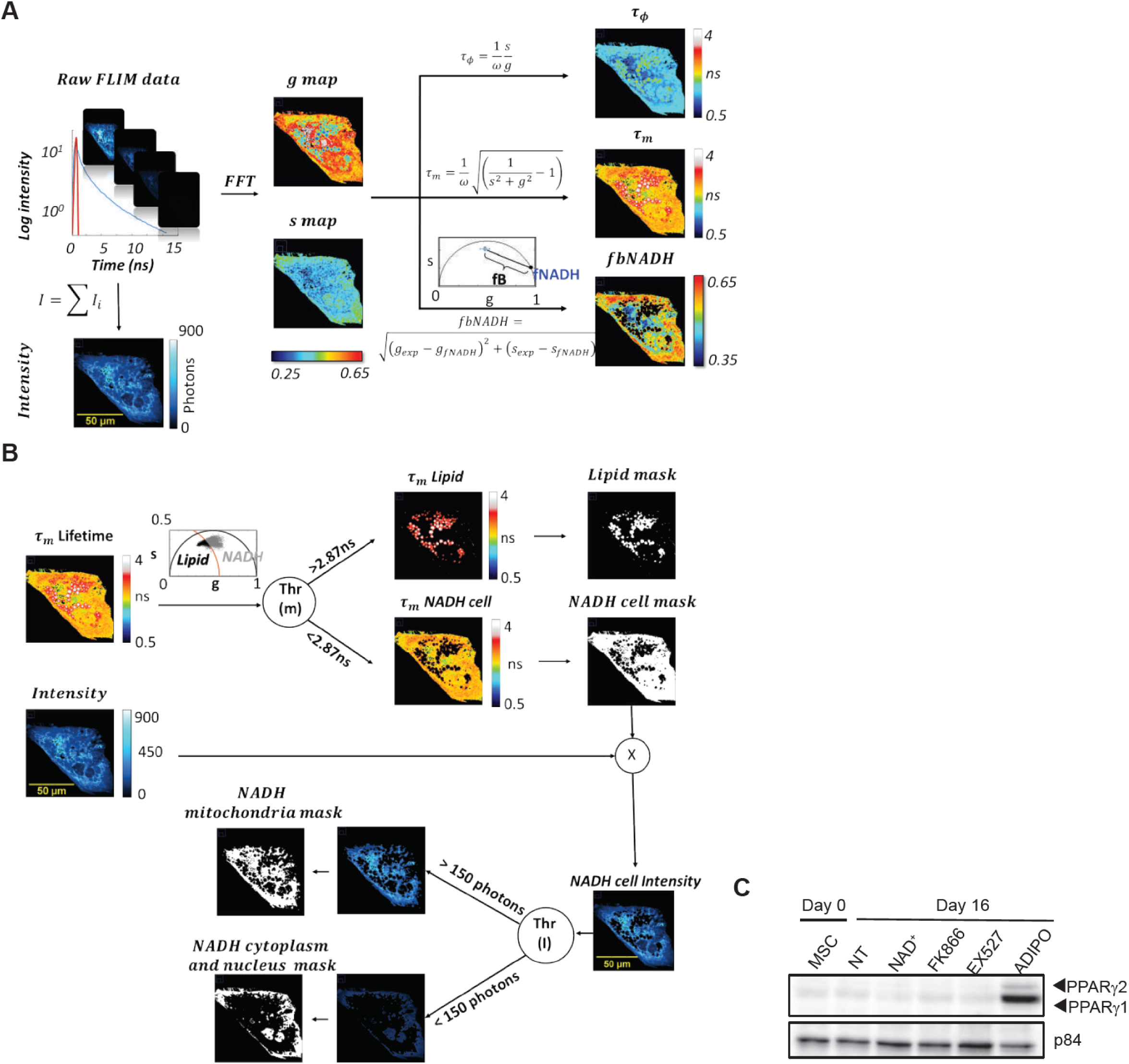
Image processing workflow for FLIM and Lipid and NADH segmentation. **A)** Workflow of FFT based Phasor analysis of Fluorescence Lifetime Microscopy images. **B)** Workflow of sub-cellular segmentation based on lifetime and intensity thresholds. A threshold on modulation lifetime (THR (m) = 2.87ns) is applied to separate lipid droplets and NADH in entire cell. The pixels of the FLIM image with τm > THR (m) (black points in the phasor plot) are assigned to lipid droplets while the pixels with τm < THR (m) (grey points in the phasor plot) are assigned to the NADH signal in the rest of the cell. A threshold (150 photons) on intensity of cell NADH is applied to segment mitochondria and nucleus plus cytoplasm. The pixels with intensity> THR (I) are assigned to mitochondria while pixels with Intensity< THR (I) are assigned to the cytoplasm and nucleus. **C)** PPARγ1 and PPARγ2 protein expression levels were measured by western blot in whole cell extracts from hMSC untreated (NT) or treated with the indicated compounds for 16 days. Terminally differentiated adipocytes (ADIPO) were also included. p84 was used as loading control.

**Fig. S2.**
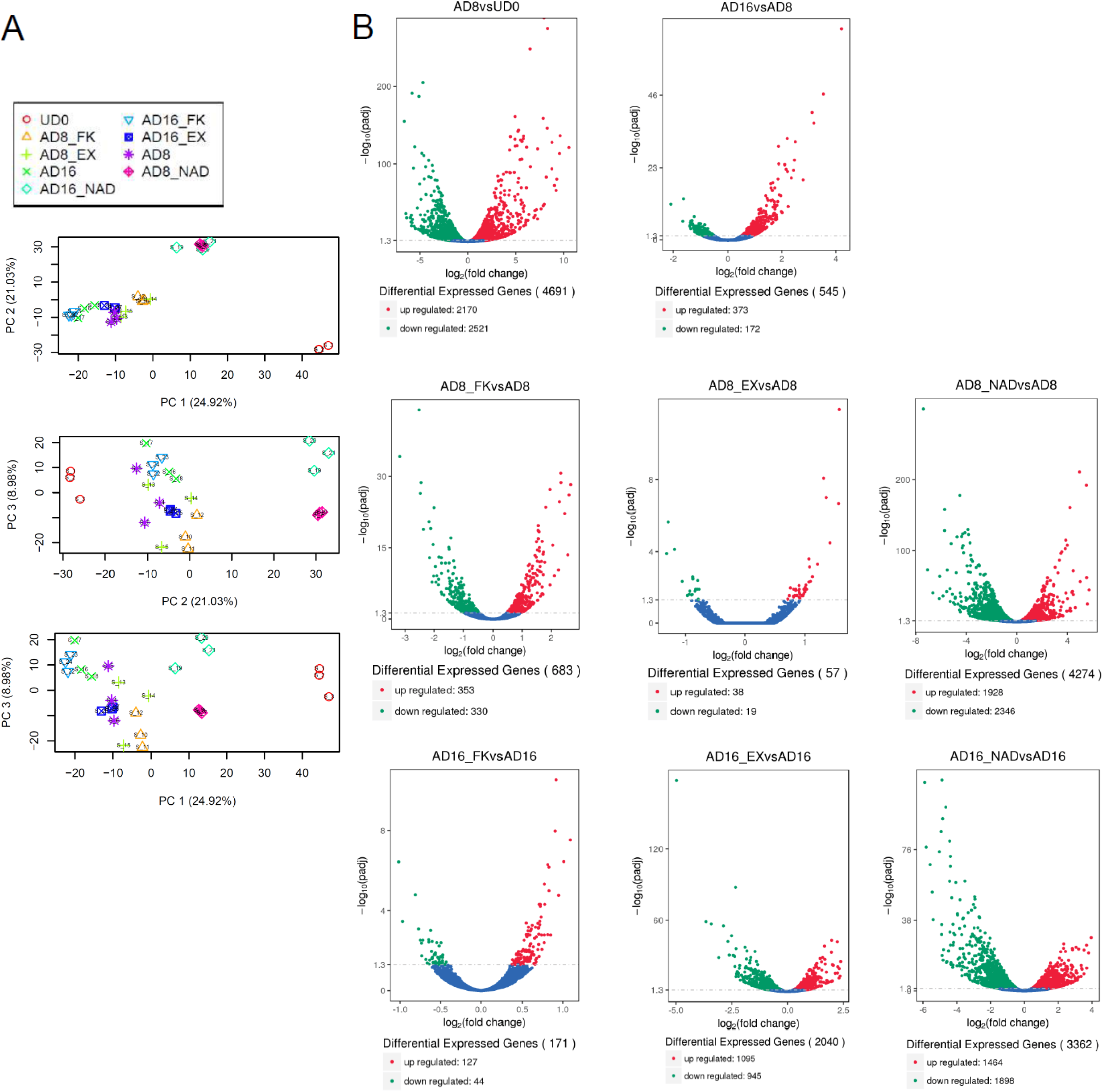
RNA-seq data analyses. **A)** Principal component analyses (PCA) was computed and plotted in two dimensional PCA score plots, showing clustering of undifferentiated hMSC (UD) vs differentiated cells (top), NAD+-treated (AD8_NAD, AD16_NAD) vs untreated cells (middle) and day 8 (adipocyte commitment) vs day 16 (terminally differentiated adipocytes) (bottom). **B)** Volcano plots show DE genes from the indicated comparisons (FDR-adjusted P-value <0.05).

**Fig. S3.**
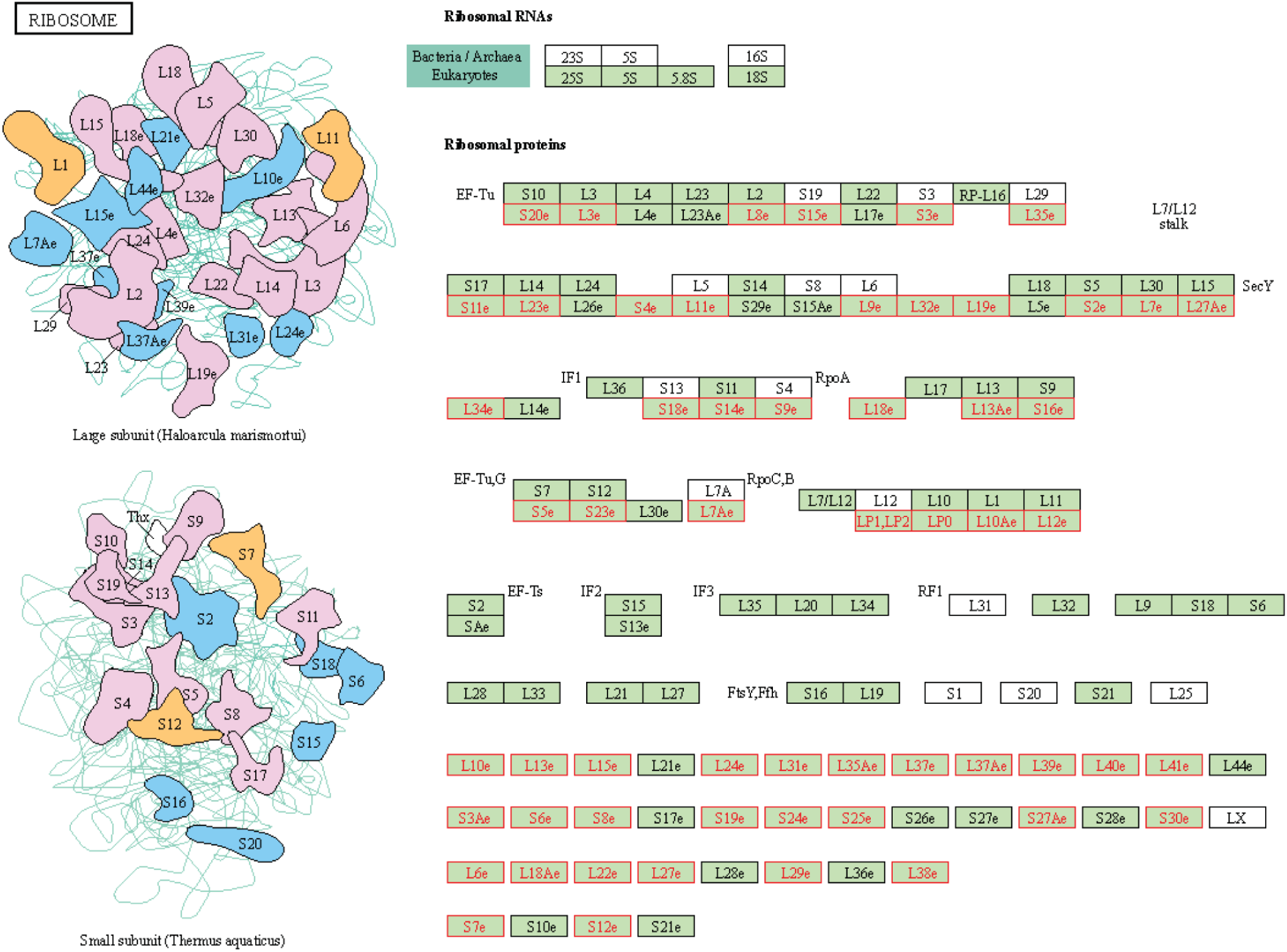
Expression of the ribosomal pathway is impaired by excess of NAD+ during adipogenesis. Illustration of the Ribosomal Pathway according to the KEGG. Ribosomal components are illustrated (left) and listed (right). Ribosomal proteins whose mRNA expression is consistently downregulated by NAD+ treatment during adipogenesis are highlighted in red color..

**Fig. S4.**
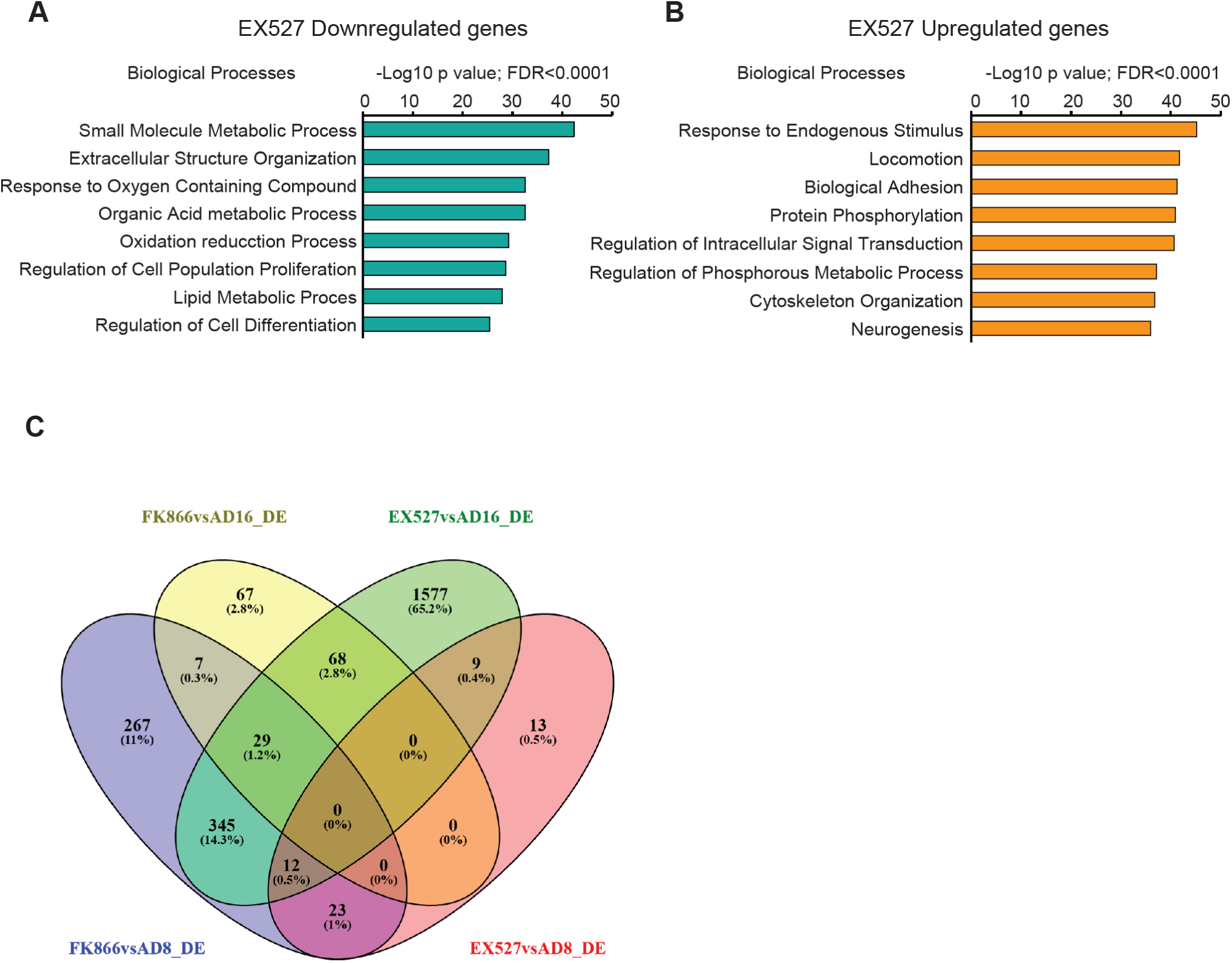
Transcriptional rewiring during adipogenic differentiation triggered by SIRT1 or NAPT inhibition. **A**, **B**) Biological processes enrichment analyses from genes downregulated (**A**) or upregulated (**B**) by EX527 treatment during adipogenic differentiation, at day 16 after induction, compared with untreated, terminally differentiated adipocytes. C) Venn diagram shows overlapping DE genes between indicated comparisons: FK866vsAD8_DE and FK866vsAD16_DE: mRNA was analyzed from cells during adipogenic differentiation (day 8 or day 16) from untreated (AD) or treated with 1nm FK866 during differentiation. EX527vsAD8_DE and EX527vsAD16_DE: mRNA was analyzed from cells during adipogenic differentiation (day 8 or day 16) from untreated (AD) or treated with 50μM EX527 during differentiation.

**Fig. S5.**
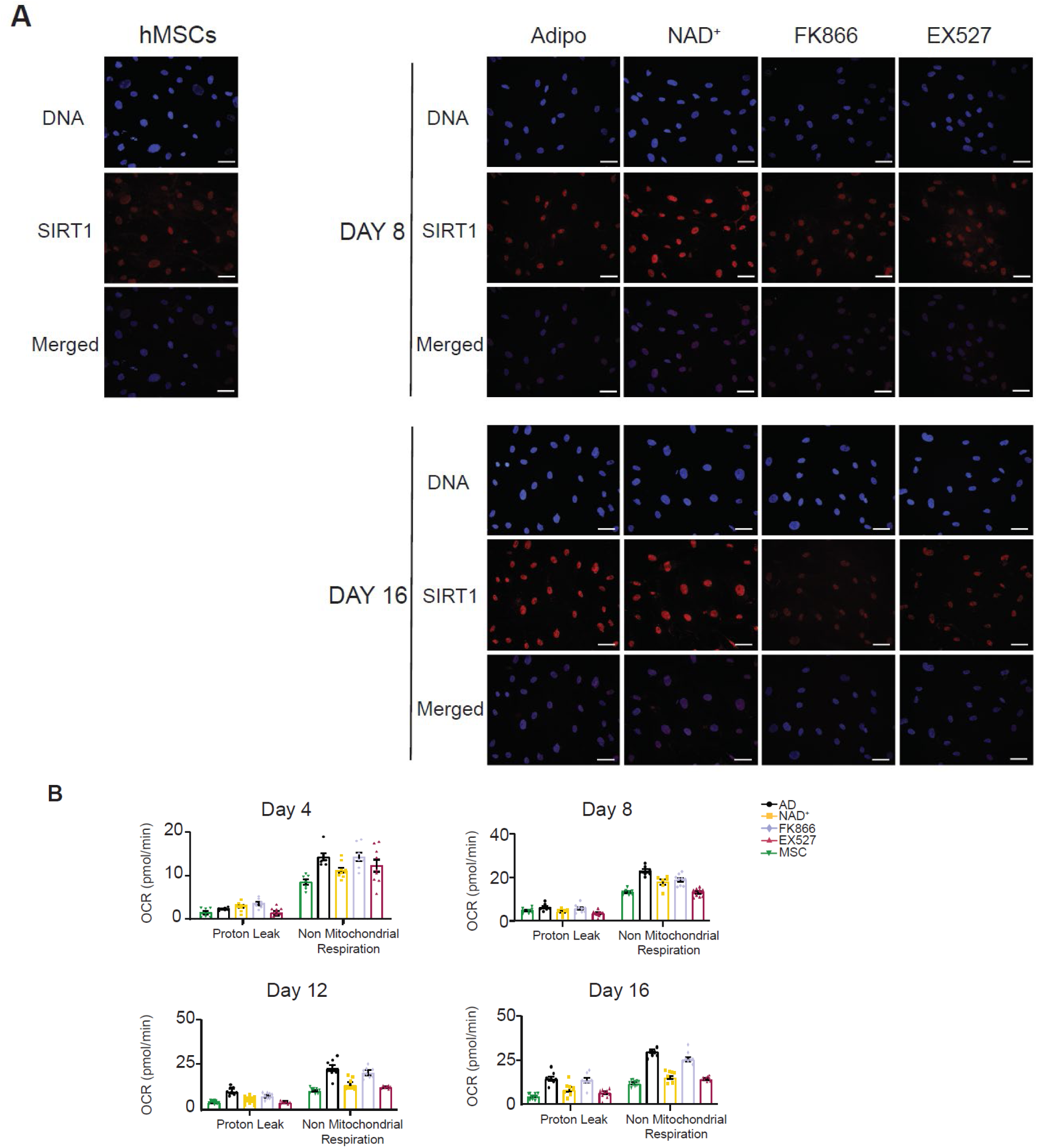
SIRT1 levels and mitochondrial bioenergetics during adipogenic differentiation. **A)** SIRT1 protein levels and subcellular localization were analyzed by immunofluorescence at days 8 and 16 after adipogenic induction on hMSC. Cells were either untreated (Adipo), or treated with the indicated compounds. n= 2 biological and 7 technical replicates. **B)** Mitochondrial bioenergetic parameters calculated from extracellular flux analyses: Proton leak and non-mitochondrial respiration. AD: adipogenic induced cells; NAD^+^ adipogenic induced cells treated with 5 mM NAD+; FK866: adipogenic induced cells treated with 1 nM FK866; EX527: adipogenic induced cells treated with 50 μM EX527. MSC: untreated, undifferentiated hMSC.

**Fig. S6.**
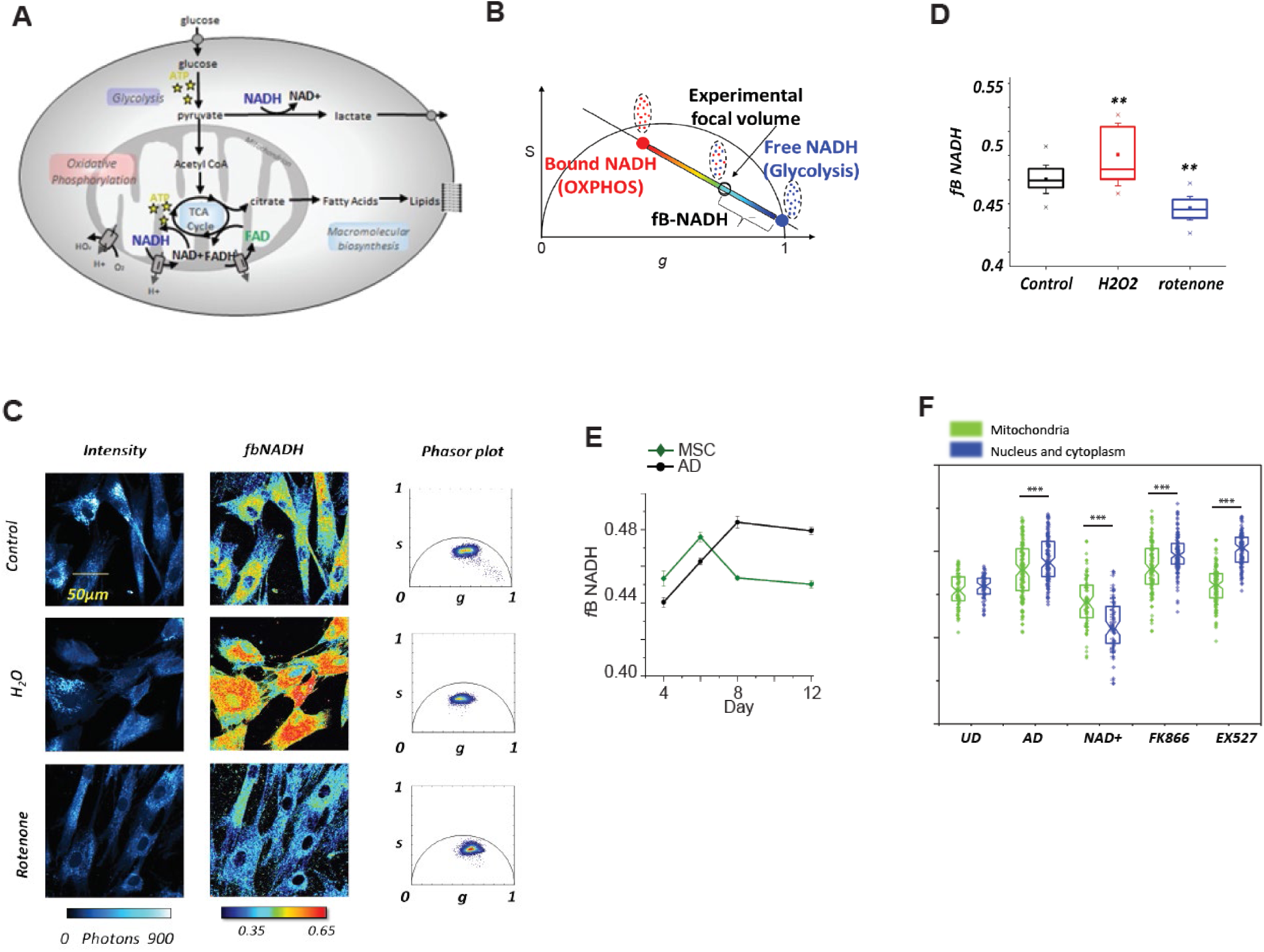
Metabolic trajectories assessed by 2P-FLIM on NADH. **A)** Schematic representation of cellular metabolism. Glucose breakdown through glycolysis and the TCA cycle generates reduced NADH and FADH2. Quiescent cells have a basal rate of glycolysis, converting glucose to pyruvate, which is then oxidized in the TCA cycle. As a result, the majority of ATP is generated by oxidative phosphorylation (OXPHOS). Non proliferating, differentiated cells are characterized by a low NADH/NAD^+^ ratio. During proliferation, the large increase in glycolytic flux rapidly generates ATP in the cytoplasm. Most of the resulting pyruvate is converted into lactate by lactate dehydrogenase A, which regenerates NAD^+^ from NADH. Proliferating cells are characterized by a high NADH/NAD^+^ ratio. Rotenone blocks the respiratory chain via complex I while H_2_O_2_ increase the NAD^+^:NADH ratio. **B)** Metabolic trajectory between free NADH and bound NADH indicates a shift from a glycolytic to a OXPHOS cellular phenotype as free/bound NADH ratio corresponds to NAD^+^:NADH ratio. The fraction of bound NADH (fB_NADH) of the experimental point is graphically calculated from the location of free NADH. **C)** Representative images of intensity, fB_NADH and phasor plot of hMSCs with different treatments: control, rotenone (respiratory chain inhibitor) and H2O2 (induces oxidative stress). Accumulation of reduced NADH by blocking the respiration chain shifts the cellular metabolic signature toward free NADH, while oxidative stress shifts the cellular metabolic signature towards bound NADH. **D)** Quantification of fraction of bound NADH in a ROI with different metabolic treatments. One-way ANOVA followed by Tukeýs post test. * p <0.05, ** p <0.01. **E)** Quantification of fraction of bound NADH during adipogenic differentiation at with (black) or without adipogenic culture medium (dark green). Data is presented as mean ±SEM F) Quantification of fB_NADH in mitochondria (green) and in nucleus/cytoplasm (blue) in single cells at day 8 of adipogenic differentiation in the absence (AD) or in the presence of the indicated treatments. hMSC (UD) were also assessed. n= 63-125 cells; ***p< 0,001, Student’s t-test.

**Fig. S6.**
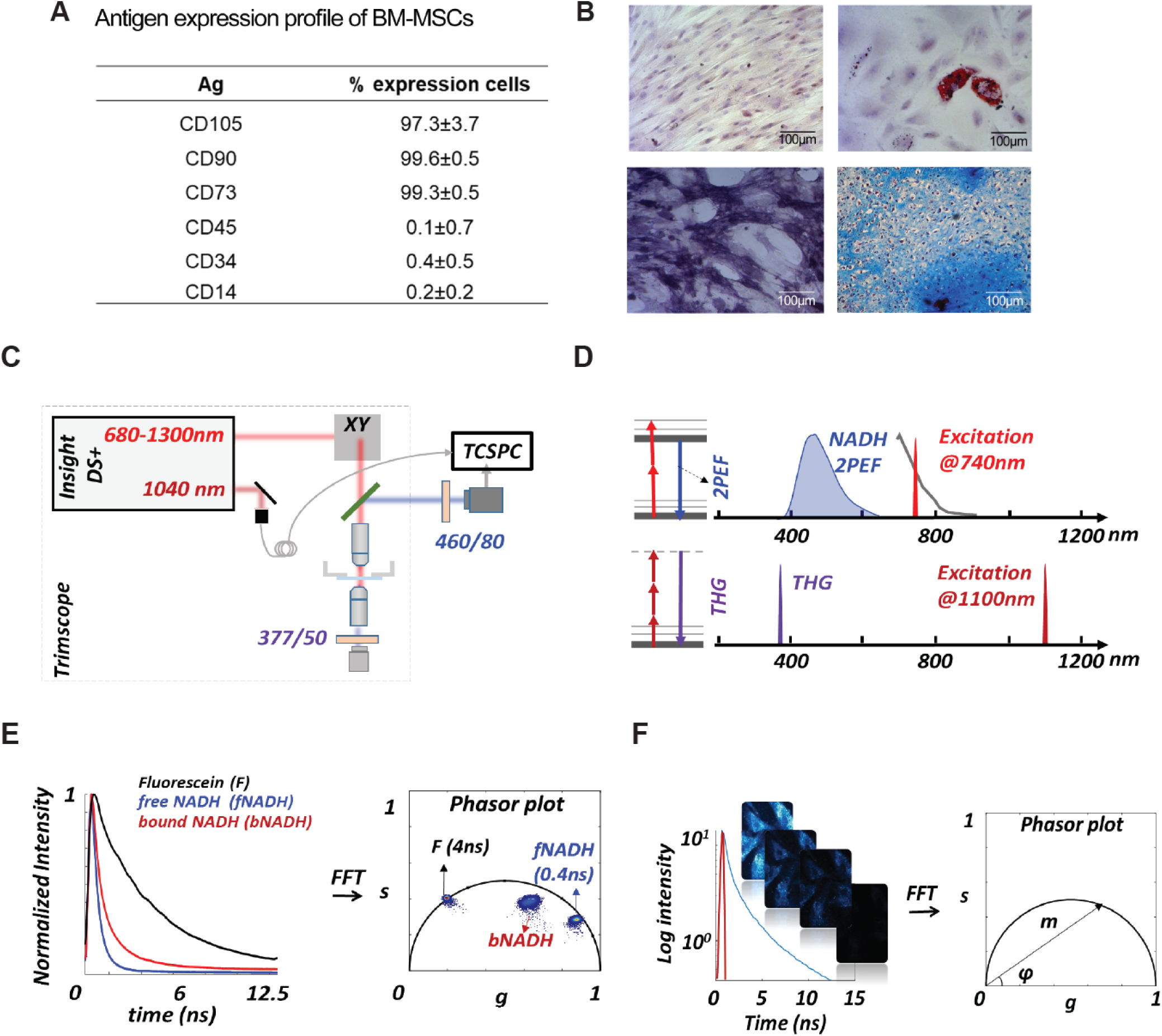
hMSC characterization and experimental setup. **A)** Expression cell markers in MSCs was determinated by Flow Cytometry, data correspond to mean percentage of cells positive to each marker ± SD, n=3 biological replicates. **B)** hMSCs stained with toluidine blue (top left), adipogenic differentiation was determined by the presence of lipid vacuoles positive to ORO (top right), osteogenic differentiation was determined by Alkaline Phosphatase assay (bottom left) and chondrogenic differentiation was assessed by matrix positive to Alcian blue in cryosections of micromasses (bottom right). n=3 biological replicates. **C)** Scheme of the experimental setup used for this work. **D)** Principles of 2 photon excitation fluorescence and THG signal generation: Two-photon excitation of NADH occurs at 740 nm with emission collected with band-pass filters centered at 460nm, which resulted primarily from NADH. The excitation of THG occurred at 1100nm, with emission collected with a band-pass filter centered at 377nm. **E)** Example of fluorescence intensity decay of fluorescein and free and bound NADH in solution and their locations in the phasor plot. **F)** The multi-exponential fluorescence intensity decay in every pixel of the image is transformed with a Fourier transform; the real (g) and imaginary (s) parts are plotted in the graphical phasor plot.

**Dataset S1 (separate file). Summary of mapping results.** Sample name: sample identification; Total reads: total clean reads suitable for analysis; Total mapped: numbers of reads being mapped on the genome; Uniquely mapped reads: numbers of reads being mapped to a single position of the genome; Multiple mapped reads: numbers of reads being mapped to more that one genomic sites; Total mapping rate: (mapped reads)/(total reads)*100; Uniquely mapping rate: (uniquely mapped reads)/(total reads)*100; Multiple mapping rate: (multiple mapped reads)/(total reads)*100.

**Dataset S2 (separate file).** List of differentially expressed genes from NAD+-treated cells genes and their functional analyses.

**Dataset S3 (separate file):** List of DE genes in from EX527-treated cells at terminal differentiation (day 16) and their functional analyses.

**Dataset S4 (separate file):** List of DE genes in from FK866-treated cells at terminal differentiation (day 16) and their functional analyses

## Notes

### Competing Interest Statement

The authors have declared no competing interest.

